# Activation mechanism of a small prototypic Rec-GGDEF diguanylate cyclase

**DOI:** 10.1101/2020.08.28.271692

**Authors:** Raphael D. Teixeira, Fabian Holzschuh, Tilman Schirmer

**Affiliations:** Structural Biology, Biozentrum, University of Basel, Klingelbergstrasse 70, 4056 Basel, Switzerland

## Abstract

Diguanylate cyclases (DGCs) synthesising the bacterial second messenger c-di-GMP are found to be regulated by a variety of sensory input domains that control the activity of their catalytical GGDEF domain. As part of two-component systems, they are activated by cognate histidine kinases that phosphorylate their Rec input domains. DgcR from *Leptospira biflexa* is a constitutively dimeric prototype of this class of DGCs. Full-length crystal structures revealed that BeF_3^-^_ pseudo-phosphorylation induces a relative rotation of two rigid halves in the Rec domain. This is coupled to a reorganisation of the dimeric structure with concomitant switching of the coiled-coil linker to an alternative heptad register. Finally, the activated register allows the two substrate-loaded GGDEF domains, which are linked to the end of the coiled-coil *via* a localised hinge, to move into a catalytically competent dimeric arrangement. Bioinformatic analyses suggest that the binary register switch mechanism is utilised by many DGCs with N-terminal coiled-coil linkers.

## Introduction

C-di-GMP is a near-ubiquitous bacterial second messenger responsible for the coordination of a variety of cellular processes and behaviour, including motility, biofilm formation, virulence and cell cycle progression (Jenal et al., 2017). Intracellular levels of c-di-GMP are regulated by the opposing actions of diguanylate cyclases (DGCs), which contains a GGDEF domain and synthetize c-di-GMP, and phosphodiesterases (PDEs), responsible for the degradation of this second messenger via an EAL or HD-GYP domain (Simm et al., 2004). Not unfrequently, dozens of these enzymes can be encoded by one single genome with each of the proteins containing distinct sensory input domains that can sense/receive diverse signals like O_2_, light and metals (Tarnawski et al., 2015)(Glantz et al., 2016) (Zähringer et al., 2013). This allows bacteria to detect intracellular and environmental cues and respond promptly by adjusting c-di-GMP levels which will then be detected by specific receptors. Common input domains are GAF and PAS, which can recognise a variety of molecules, and response regulator receiver domains (Rec), which as part of two component systems are phosphorylated by cognate histidine kinases (HKs) (Zoraghi et al., 2004) (Henry and Crosson, 2011) (Gao and Stock, 2010).

DGCs catalyse the condensation of two molecules of GTP to yield the 2-fold symmetric c-di-GMP product. This requires the juxtaposition of two GTP loaded DGC domains in appropriate 2-fold related arrangement to form a catalytically competent GGDEF dimer that enables nucleophilic attack of the deprotonated O3’ hydroxyl onto the phosphorous of the other GTP molecule (Schirmer, 2016). The first characterized full-length DGCs were PleD and WspR of Rec – Rec – GGDEF and Rec - GGDEF domain organization, respectively, which were also studied in the beryllofluoride (BeF_3^-^_) modified form known to mimic phosphorylation (Wemmer and Kern, 2005). It was shown that, upon this modification, PleD shifts from a monomer to a catalytically active dimer (Wassmann et al., 2007), whereas the behaviour of WspR was more complex in that it enhanced tetramer formation (De et al., 2009). Later, structural and biochemical analyses on DGCs with other input domains revealed that these enzymes can exist also as constitutive dimers. Zinc binds to the CZB domain of DgcZ and prevents productive encounter of the GGDEF domains by restraining domain mobility (Zähringer et al., 2013). DosC has a globin domain with bound heme to sense oxygen (Tarnawski et al., 2015), and lastly, the bacteriophytochrome PadC senses red light through its PHY domain to activate the GGDEF domain (Gourinchas et al., 2017).

Almost invariably, input and catalytic domains of DGCs are connected by a dimeric coiled-coil that can vary in length. We proposed earlier that the constituting helices could change their crossing angle and/or azimuthal orientation to allow or prevent productive encounter of the two GGDEF domains (chopstick model, (Schirmer, 2016)). This mechanism would be a generalization of the scissors with fixed pivot blades model ascribed to histidine kinases signalling (Lowe et al., 2012). To test for the mechanism, we selected a prototypical minimal DGC with known input signal. LEPBI_RS18680 from *Leptospira biflexa*, hereafter called DgcR (**d**i**g**uanylate-**c**yclase controlled by **R**ec), is a Rec - GGDEF protein with a short domain linker (Fig. 1a), thus making this enzyme attractive for studying its conformational states by crystallography.

**Figure 1.**
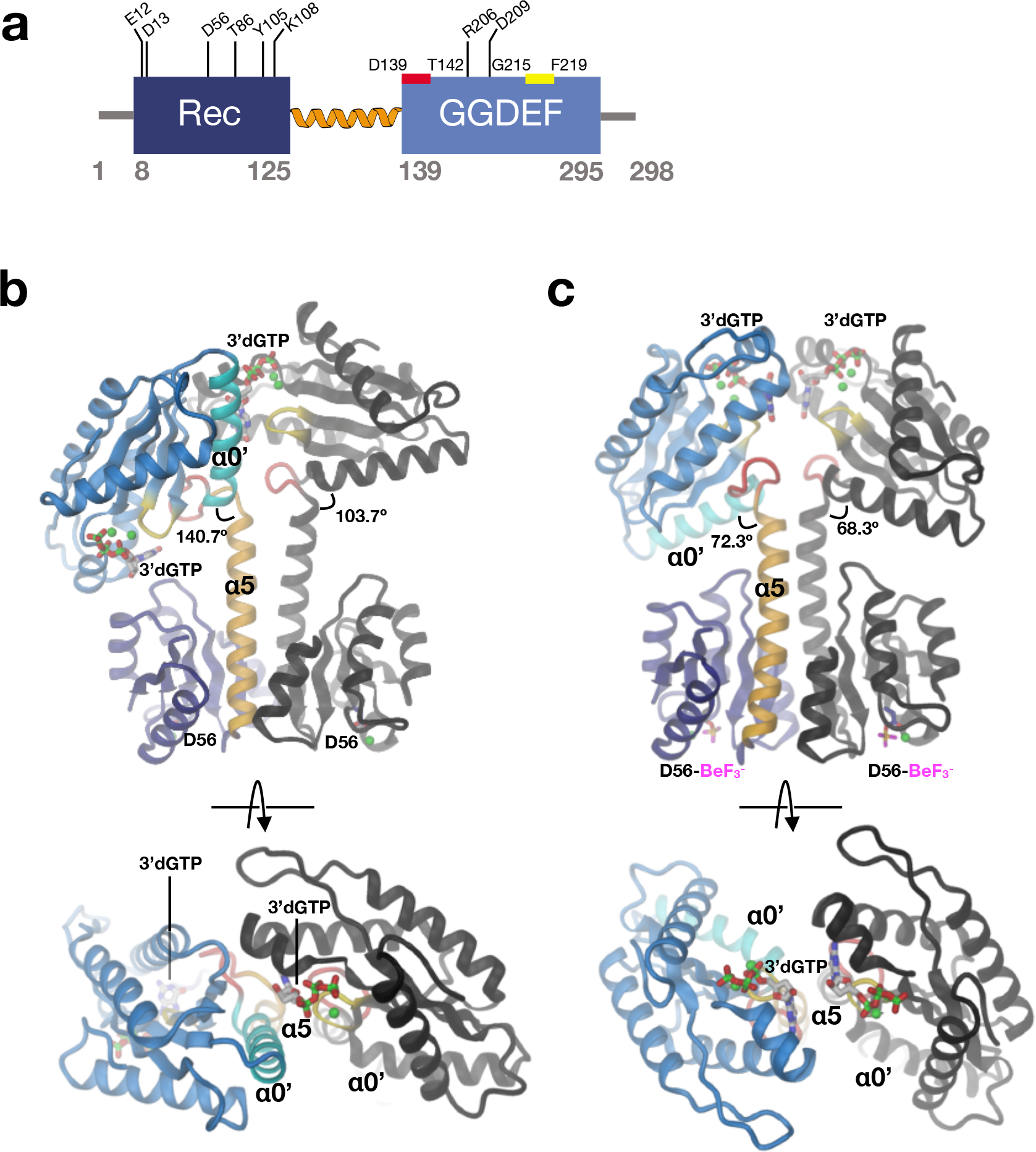
Native and activated DgcR dimers adopt distinct domain arrangements. **a)** DgcR domain organisation with important features and residues highlighted. The Rec and the GGDEF domains are linked by the extension (orange) of the C-terminal Rec helix. Red and yellow bars indicate the DxLT and GGDEF motif, respectively. Note the presence of the RxxD inhibition site. **b-c)** Side and top views of DgcR_nat (**c**) and DgcR_act (**d**) dimers. Within one protomer, domains and important elements are highlighted by color. The C-terminal Rec helix (α5) is colored in gold, the wide turn in red, the N-terminal GGDEF helix (α0’) in cyan and the characteristic β-hairpin (with GGEEF sequence) in yellow. The BeF_3_^-^ modified aspartates of the Rec domains, the bound Mg^++^ ions (green), and the 3’dGTP substrate analogues bound to the GGDEF active sites are shown in full. In both cases, the Rec domains obey 2-fold symmetry. The GGDEF domains are related by 2-fold symmetry (with the two 3’dGTP ligands opposing each other) only in DgcR_act, while in DgcR_nat they are related by approxmately 90 °.

*Leptospira* is a bacterial genus composed of more than 30 species, among them some pathogenic representatives responsible for causing leptospirosis, a worldwide zoonosis that affects more than one million people and accounts for 60,000 deaths per year (Karpagam and Ganesh, 2020). *Leptospira biflexa* is a saprophytic species used as a model to study *Leptospira* biology (Pětrošová and Picardeau, 2014). It contains an additional extra-chromosomal element of 74 kb (p74) that codes for 56 proteins including DgcR. DgcR was also chosen, because it shares the same domain organization and linker length as Rrp1 from *Borrelia burgdorferi*, the pathogen responsible for causing the Lyme disease. Rrp1 is the only DGC encoded by *B. burgdorferi* genome and is essential for bacterial survival in the tick host (He et al., 2011).

Here we show by biophysical and crystallographic analyses that DgcR is a constitutive dimer that changes coiled-coil geometry and domain arrangement upon pseudo-phosphorylation. The chopstick model is generally confirmed, but, upon activation, DgcR shows an unexpected translational register shift. Bioinformatic analyses suggest that the observed activation mechanism is most likely operational in most diguanylate cyclases of Rec – GGDEF organisation, but also in some other DGCs.

## Results and discussion

### DgcR is a constitutive dimer that gets activated by domain rearrangements

To reveal the structural changes accompanying the activation of Rec - GGDEF DGCs we determined the full-length crystal structures of DgcR in native and pseudo-phosphorylated (BeF_3^-^_ modified) state. A DgcR variant (R206A/D209A, abbreviated DgcR_AxxA) that had the putative allosteric inhibition site (Fig. 1a) mutated was used to avoid locking the enzyme in a product inhibited conformation. Crystallization was performed in presence of 3’-deoxy-GTP (3’dGTP), which is a non-competent substrate analogue due to the absence of the 3’-hydroxyl group.

The structure of native DgcR_AxxA (called DgcR_nat) was solved by molecular replacement to 2.2 Å resolution. There is one dimer in the asymmetric unit with the protomers held together by extensive isologous contacts between the Rec domains involving their α4 - β5 - α5 face (Fig. 1b). The Rec domain shows the canonical (βα)5 fold (rmsd of 1.5 Å for 116 Cα atoms with respect to PhoP, 2PKX), but with the C-terminal α5 helix considerably extended and forming together with its symmetry mate a coiled-coil leading to the GGDEF domains. A Mg2+ ion is bound to the acidic pocket formed by E12, D13, and the phosphorylatable D56.

The structure of the GGDEF domain is very similar to others in the PDB database (rmsd of 1.4 Å for 157 Cα atoms with respect to PleD, 2V0N) and shows the canonical (β1-α1-α2-β2-β3-α3-β4-α4-β5) topology of nucleotidyl cyclases of group III (Sinha and Sprang, 2006) with an N-terminal extension that starts with a characteristic wide turn showing a DxLT motif followed by helix α0 that leads to β1 (Fig. 1b, see also (Schirmer, 2016). The GG(D/E)EF motif is located at the turn of the β2-β3 hairpin. Again as observed in other structures (Wassmann et al., 2007), the guanine base of the substrate analogue is bound to a pocket between α1 and α2 and forms H-bonds with N182 and D191, whereas the two terminal phosphates are H-bonded to main chain amides of the short loop between β1 and α1.

Additionally, the γ-phosphate forms ionic interactions with K289 and R293. Two magnesium ions are bound to the usual positions being complexed to the β- and γ-phosphates and the side-chain carboxylates of D174, E217, and E218.

The GGDEF domains do not obey the 2-fold symmetry of the Rec domains, but form a relative angle of about 90°. Thus, the two active sites with the bound GTP analogues do not face each other rendering this constellation clearly non-productive. Though the constellation may be determined to some extent by crystal packing, it demonstrates considerable inter-domain flexibility. Comparison of the main-chain torsion angles reveals that the relative rotation can be traced back to a 169° change in a single torsion, namely around the Cα - C bond of residue 136 (ψ136, Fig 2a). Thus, the hinge locates to the C-terminal end of Rec α5, with the following residue I137 being packed against the Y149 from the end of the GGDEF α0’ helix in both chains (Figs. 2b and c). As noted before (Schirmer, 2016), the conserved residue N146 (see sequence logo in Fig. 2 - figure supplement 1) is capping both α5 and α0’, but only in the A-chain.

**Figure 2.**
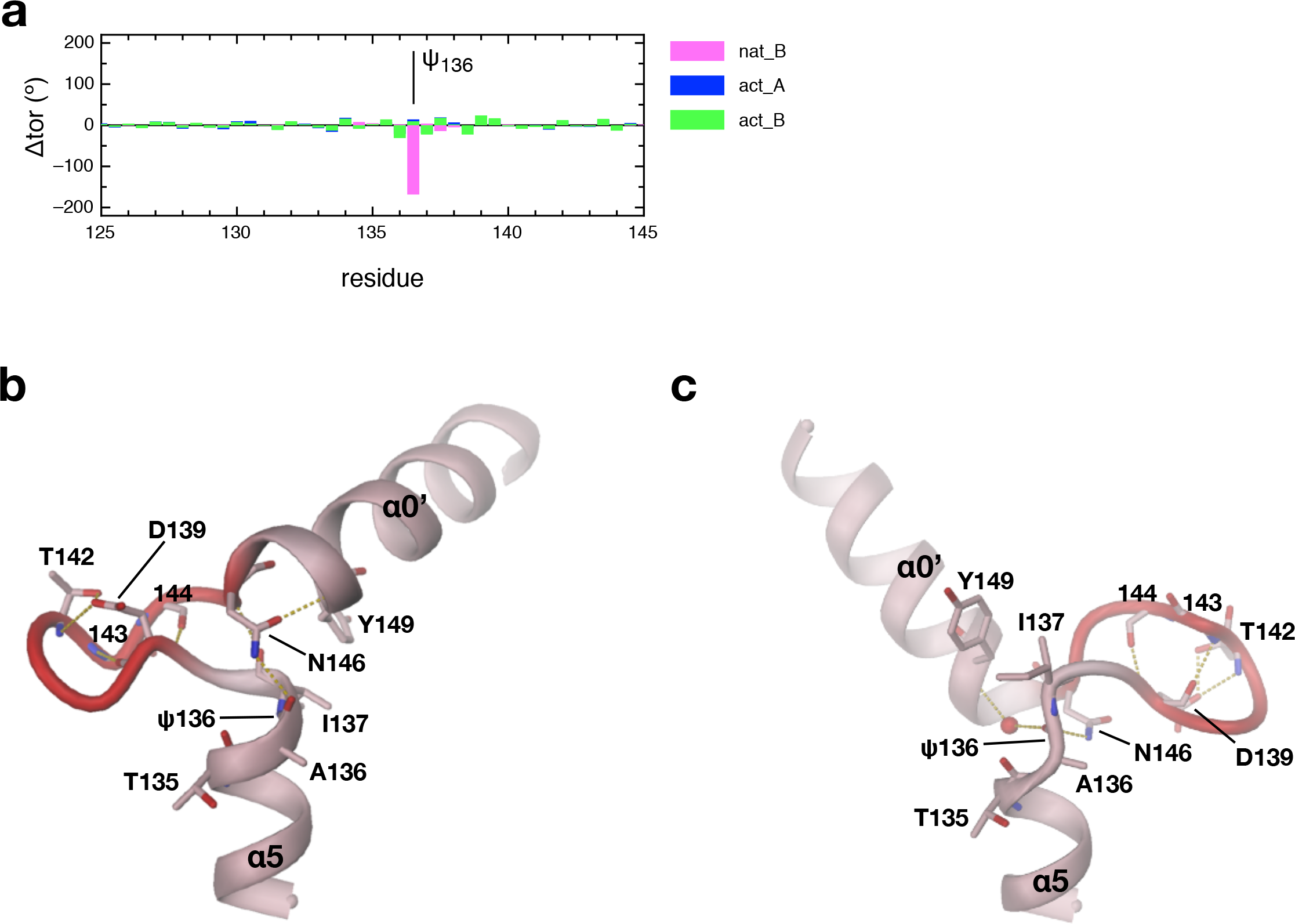
Inter-domain hinge revealed by comparison of the two DgcR_nat protomers. **a)** Backbone torsion angle comparison of DgcR_nat and DgcR_act (nat_B, act_A and act_B) relative to nat_A. Note that the two DgcR_nat chains show a localized drastic difference in the main-chain torsion angle ψ136. **b-c)** Detailed view of the inter-domain hinge segment of chain A (panel **b**) and B (**c**) of DgcR_nat. The wide turn with the D139xLT142 motif is high-lighted in red. The 169° rotation of ψ136 drastically changes the relative angle between α5 of the Rec and α0’ of the GGDEF domain.

The structure of activated DgcR_AxxA (called DgcR_act) obtained by BeF_3^-^_ modification was solved by molecular replacement to 2.8 Å resolution (Fig. 1c). There are two dimers in the asymmetric unit that show virtually the same Rec dimer structure, but slightly different α5-helix bending and GGDEF orientations (Figure 1—figure supplement 1). As in DgcR_nat, the dimer is formed by isologous contacts between the α4-β5-α5 Rec faces and the extension of α5 forms a coiled-coil, but with an altered relative disposition, which will be described in detail further below. D56 is found fully modified by BeF_3^-^_ and its immediate environment is different compared to DgcR_nat as will be discussed in detail hereafter. The GGDEF domains are arranged symmetrical with the two bound 3’dGTP ligands facing each other, but too distant for catalysis (Fig. 1c, bottom). The GGDEF orientation relative to the Rec domain is similar as in the A-chain of DgcR_nat.

Consistent with the crystal structures and the presence of the coiled-coil in both states, in solution, DgcR is a constitutive dimer as measured by MALS both in the native and the activated form (Figure 1—figure supplement 2). Addition of substrate analogue or product was not changing this quaternary state.

### Aspartate modification induces a relative rigid-body rotation within the Rec domain

Comparison of DgcR_act with DgcR_nat (Fig. 3a) shows that, on activation, the hydroxyl group of T85 is moved towards the BeF_3^-^_ moiety to form an H-bond. The void left by this movement is claimed by Y105 that undergoes a small side-chain rotation, but does not change its rotamer. Furthermore, K108 forms ionic interactions with BeF_3^-^_ and E12 in the activated structure (Fig. 3a). In the native state, a magnesium ion is bound loosely to E13 and D56, whereas, in the active state, it is additionally coordinated by the BeF_3^-^_ moiety.

**Figure 3.**
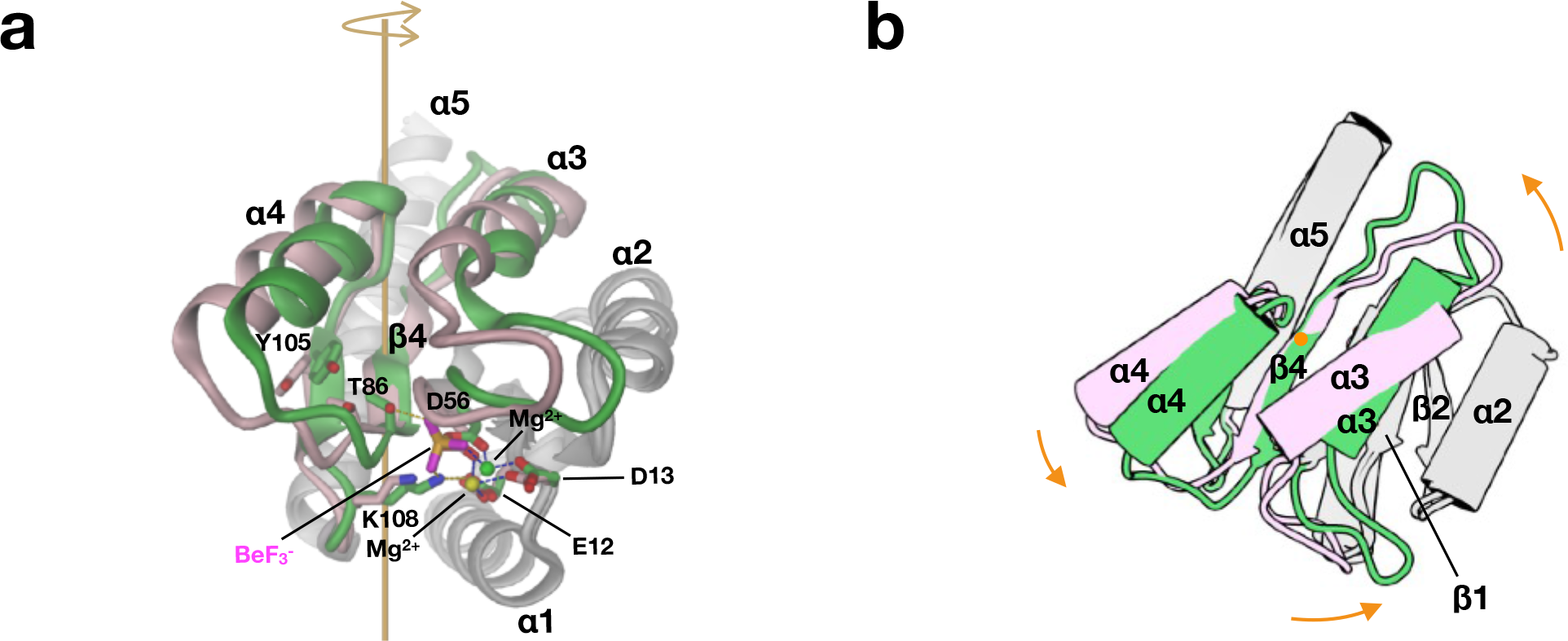
Beryllofluoride modification induces a relative 16° rotation of two Rec halves. Rigid-body 1 composed of α3, β4, α4, β5 (shown in pink and green for DgcR_nat and DgcR_act, respectively) is rotated with respect to the rest (rigid-body 2, grey) as seen after super-postion of the two grey substructures. **a)** Native and activated Rec domains with important residues shown in full viewed perpendicularly to the rotation axis of the relative rotation (orange). Tin DgcR_act, the beryllofluoride group forms an H-bond with T86 and an ionic interaction with K108 of DgcR_act. **b)** Same as **a**), but in cartoon representation and viewed along rotation axis.

Activation of DgcR is accompanied with a change in the backbone structure as identified by a DynDom analysis (Girdlestone and Hayward, 2015). The Rec domain can be divided into two parts that undergo a relative 16° rotation as shown in Fig. 3b. Thereby secondary structure elements α3 to β5 (residues 54 to 108) behave as one rigid body (rmsd = 0.83 Å /49) that rotates relative to the rest (8-53, 109-135) that superimposes with an rmsd of 1.19 Å for 67 Cα positions. Figure 3b shows that the rotation axis passes roughly perpendicular to the β-sheet through the centre of β4 (L83). Note that the phosphorylatable D56 is close to the junction between the two rigid bodies and that its Cα position only changes slightly during the transition. T86, however, with its distance of 7.5 Å from the rotation axis moves by 2.0 (Cα) to 3.3 Å (Oγ) and the motion is most pronounced (5.2 Å) for the N-terminus of α3 (P91) with its distance of about 15 Å from rotation axis. Thus, the rigid body motion changes significantly the arrangement of α4 with respect to α5, which has a profound effect on the packing of the Rec domains in the dimer.

For many Rec domains, a Y-T coupling mechanism has been described, where, upon (pseudo-) phosphorylation a threonine/serine (T86 in DgcR) is dragged towards the phosphate and the conserved tyrosine/phenylalanine (Y105 in DgcR) follows suite with a rotameric change from *gauche*+ to *trans* (Birck et al., 1999), (Bachhawat et al., 2005), (Wassmann et al., 2007). In DgcR, the conserved tyrosine is already in *trans* conformation before activation and the T and Y move concertedly towards the beryllofluoride moiety as part of a rigid-body (α3 to β5) movement (Fig. 3).

### Rigid body rotation induces repacking of Rec domains within the dimer

In both states, the Rec domains form 2-fold symmetric dimers with the contacts mediated by isologous interactions between the α4 - β5 - α5 surfaces (Fig. 4). However, due to the rigid-body motion within the protomer and the concomitant relative displacement of α4 and α5 (Fig. 3), the association of the α4 - β5 - α5 faces is different in the native and the activated state. Therefore, the two dimers superimpose rather poorly (rmsd = 3.1 Å/119 Cα positions) with the β-sheets of the protomers showing a difference in orientation of about 15° (Fig. 4a).

**Figure 4.**
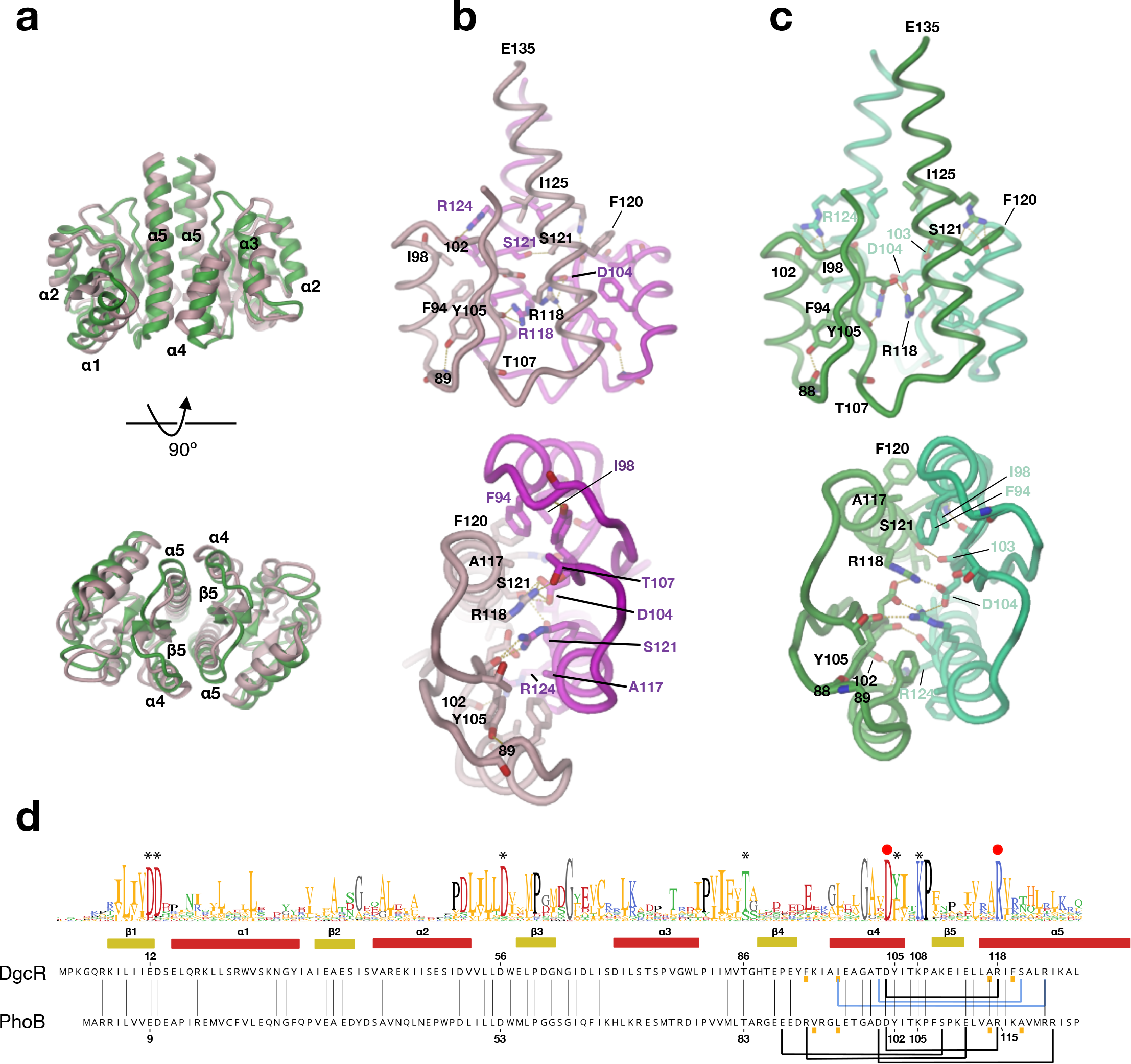
Distinct packing of Rec domains upon BeF_3^-^_ modification. **a)** Superimposed dimers with native structure in pink and activated structure in green. **b - c)** Rec dimer of DgcR_nat (**b**) and DgcR_act (**c**) with contacts between the protomers <3.2 Å in yellow. **d)** Sequence alignment of Rec domains of DgcR and PhoB with secondary structure elements of DgcR and sequence logo derived from DgcR group 1 homologs (see Fig. 10). Asterisks indicate conserved residues involved in Rec domain activation. Below the individual sequences, lines connect residues that participate in inter-domain contacts (side-chain - side-chain interactions in black, interactions involving the main-chain in blue). Red dots indicate D104 and R118 of the conserved inter-molecular salt-bridge.

The native Rec dimer (Fig. 4b) with a buried surface area of 980 Å2 is held together by an extended apolar contact of α5 (A117, F120) with α4 (F94, I98), an ionic interaction of D104 with R118, an H-bond between main-chain carbonyl 102 and R124 (both β5 - α5 contacts). All aforementioned residues are well defined with the exception of the R118 side-chain, which probably has several alternative conformations, but all placing the guanidinium group close to D104 and to its symmetry mate. Finally, and most relevant for the allosteric regulation of the C-terminal GGDEF effector domains, there are regular coiled-coil interactions across the symmetry axis between the C-terminal halves of the α5 helices starting with S121. These will be discussed in the next chapter.

The activated Rec dimer (Fig. 4c) with a buried surface area of 850 Å2 shows the same apolar α5 - α4 cluster as the native dimer, but with the residues repacked in-line with the aforementioned relative displacement of α4 and α5 within the protomer. At the centre of the interface, D104 shows a well-defined, intermolecular salt-bridge with R118, but also with R118 from the same chain. As in the native dimer, the R124 and S121 side-chains form intermolecular H-bonds, but with other partners compared to the native interactions (main-chain carbonyls of 98 and 103, respectively).

A BLAST search revealed that, apart from Rec - GGDEF orthologs, the sequence of the DgcR Rec domain is most similar to that of OmpR-like transcription factors (Fig. 4d). These have a Rec - DNA-binding domain architecture and have been shown to dimerize *via* the Rec α4 - β5 - α5 face upon activation to allow binding of their effector domains to DNA (Draughn et al., 2018). Indeed, a structure search of the DgcR_act dimer against the PDB retrieved as top hit (rmsd = 1.5 Å/228 Cα positions) the BeF_3^-^_ activated Rec domain of PhoB (1ZES) (Bachhawat et al., 2005). Most of the intermolecular interactions are thereby conserved, in particular the central salt-bridge D109 - R118 (DgcR numbering), or conservatively replaced (Fig. 4d). To our knowledge, no response regulator with a DNA binding effector domain has yet been observed as a constitutive α4 - β5 - α5 dimer (for a review, see (Gao et al., 2019), which is probably due to their small or absent coiled-coil linkers.

Summarizing, beryllofluoride-modification of D56 induces a relative rigid body motion in the Rec domain that changes the relative disposition of α4 and α5. Consequently, since both helices are part of the Rec - Rec interface, the relative arrangement of the protomers and, thus, of the two α5 helices of the dimer is changed (compare top panels of Figs. 4b and c). This change is supposed to be crucial for the allosteric regulation of the C-terminal GGDEF domains as will be discussed in the following.

### Relative translation of C-terminal Rec helices changes coiled-coil register

The DgcR Rec α5 helix is longer by about 3 turns (10 residues) compared to that of canonical Rec domains. In the dimer, these protrusions form a 2-fold symmetric coiled-coil both in the native and the activated state (Figs. 1b-c), though with distinct relative arrangement. Both constellations are stabilized by isologous contacts between predominantly hydrophobic residues that obey a heptad-repeat pattern (Figs. 5a and b). Thereby, I125 and L132 contribute to the contact in both structures (position a; persistent contacts), but with the side-chains interacting with their symmetry mates from opposite sides depending on the state (see e.g. the 132 - 132 contact in Fig. 5a). In contrast, other residues contribute either only to the native (L128, T135) or the activated (H129, A136) constellation (positions *d, e*; conditional contacts).

**Figure 5.**
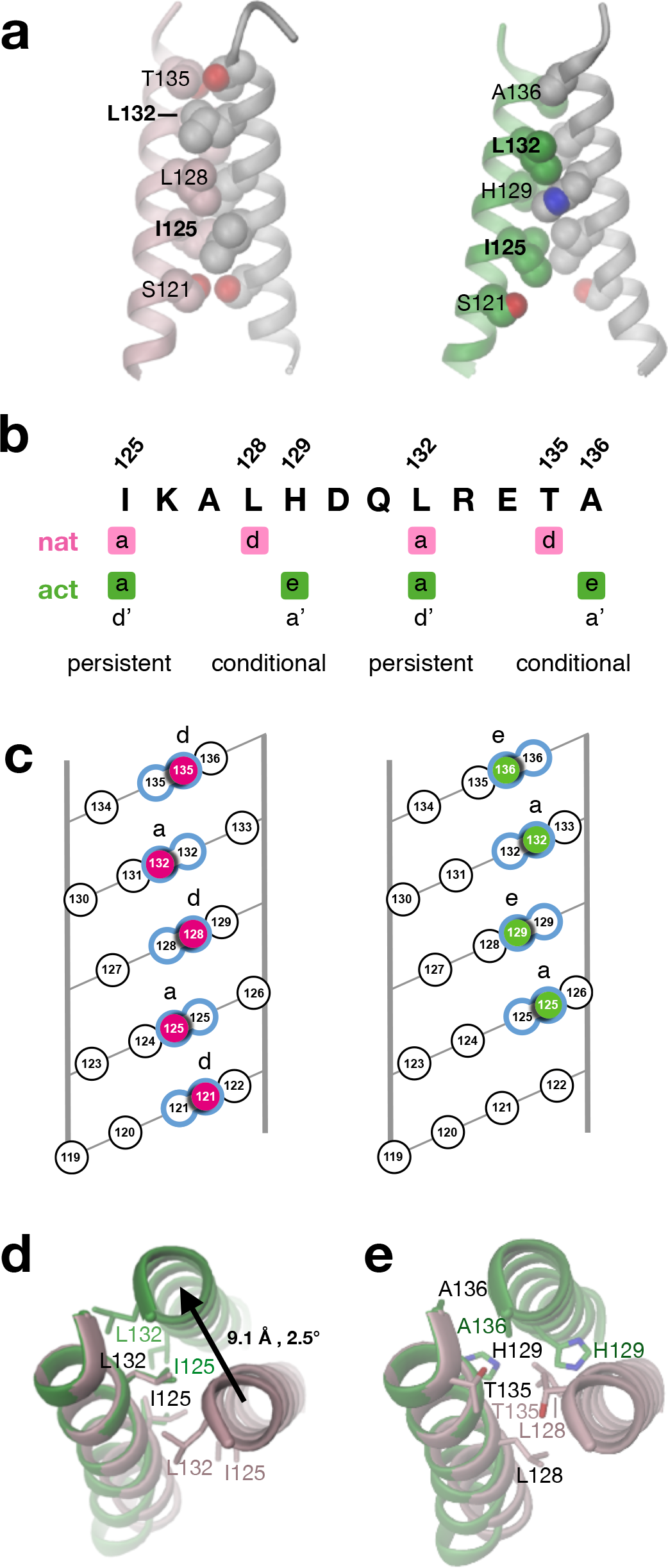
Coiled-coil linker adopts two alternative registers depending on activation state. **a)** Side view of parallel coiled-coil linker of DgcR_nat (left) and DgcR_act (right) with residues that form contacts with their symmetry equivalents in CPK representation. Only residues from the left helix are labeled. Bold labels indicate residues that are involved in both registers (persistent contacts). **b)** Two alternative heptad repeat patterns (a *d a d* and *a e a e*), which are adopted by DgcR_nat and DcR_act, respectively. Positions *a* are used in both registers (persistent contacts), whereas positions *d* or *e* are used only in the native or activated conformation (conditional contacts). Note, that the *a e a e* pattern can formally also be described by a *d*’ *a*’ *d*’ *a*’ pattern as indicated in the figure and used in (Gourinchas et al., 2017). **c)** Helical net representation (Crick, 1953) of coiled-coil interactions in DgcR_nat (left) and DgcR_act (right). Of the front helix, only interacting residues are shown (high-lighted by color). Interacting residues pairs are outlined in blue. **d-e)** Top view of coiled-coil after superposition of left helix. DgcR_nat and DgcR_act are shown in pink and green, respectively. For clarity, residues forming persistent and conditional contacts are shown in the separate panels **d** and **e**, respectively.

The two contact modes represent alternative knobs-into-holes packing as best seen in the helical net diagram of Fig. 5c suggesting a relative translation of the interacting helices. Indeed, superposition of one of the helices as in Figs. 5d and e reveals a large relative translation of about 9 Å. In other words, upon activation, the two helices do not roll over each other (which would be accompanied by a change in their azimuthal angles), but are translated with respect to each other to realise an alternative knobs-into-holes packing. Note, that for steric reasons this shift would require dissociation and reassemble of the constituting helices. Thus, the coiled-coil behaves like a binary switch that can assume two clearly defined states, i.e. two distinct registers.

Recently, an analogous transition in the coiled-coil linker of a diguanylate cyclase has been proposed for phytochrome-regulated PadC (Gourinchas et al., 2017). Indeed, the C-terminal end of the coiled-coil of the dark-state enzyme is in the same register as native DgcR with, amongst others, N518 and L525 forming (conditional) contacts (see PadC in Fig. 5 - figure supplement 1). Inspection of the linker sequence and dynamic considerations prompted the authors to propose an alternative register involving the neighbouring residues N519 and A526 for the illuminated state. Indeed, mutations designed to stabilize this second register were constitutively active and the coiled-coil was found in the active register (see PadC_mut in Fig. 5 - figure supplement 1), (Gourinchas et al., 2018). Though the structure of light-activated wild-type PadC is not known, it is very likely that PadC and DgcR use the same binary coiled-coil switch mechanism for DGC regulation, despite unrelated input domains.

WspR, another well-studied Rec - GGDEF diguanylate cyclase, also exhibits a “slippery” *axxdexx* heptad-repeat with *e* of the last repeat in position −3 with respect to the DxLT motif (Fig. 5 - figure supplement 1). Unfortunately, only product bound structures are available that reveal a non-productive, c-di-GMP cross-linked tetramer in which the coiled-coils emanating from the two Rec dimers are splayed apart at their ends (De et al., 2008). Most revealing, however, a GCN4 – GGDEF WspR with GCN4 interface residues in the active register (Fig. 5-figure supplement 1) was reported to be highly active and a corresponding structure (compact dimer) was predicted for active WspR (De et al., 2009).

Summarizing, the change in coiled-coil registration upon DgcR activation is accompanied by a substantial shift between the constituting helices that lead directly to the catalytic domains. Structural data on other DGCs are consistent with this finding.

### Small rotation around inter-domain hinge allows formation of competent GGDEF dimer

In the activated structure, the two GGDEF domains show no mutual interactions and their precise orientation appears to be determined by crystal contacts. However, the two bound 3’dGTP ligands face each other, though their distance (> 10 Å) is clearly too large for catalysis (see Fig. 1c). Having identified the CA - C main-chain bond of A136 as an inter-domain hinge (Fig. 2a), we tried, by small changes in ψ136 and adjoining main-chain dihedrals, to symmetrically move the GGDEF domains as rigid bodies into a catalytically competent arrangement (Michaelis-Menten complex). Indeed, only small torsional changes (Fig. 6a) were necessary to bring the (reconstructed) 3’-hydroxyl groups of each bound substrate in line with the scissile PA - O3A bond of the other substrate as required for catalysis (Figs. 6b and c). It should be considered, however, that the optimal arrangement of the catalytic domains depends obviously on the conformation of the bound substrates, as discussed in (Zähringer et al., 2013). In fact, comparison of the bound 3’dGTP ligands in DgcR_nat and DcrR_act shows variability in the ribose and α-phosphate orientation (Fig. 6 – figure supplement 1), probably due to the lack of strong interactions with the binding site. Here, we used the conformation as seen in DgcR_act. Scenarios with other substrate conformations were not explored, but the relative GGDEF arrangement would probably be similar considering the fixed hinges at the end of the activated Rec dimer.

**Figure 6.**
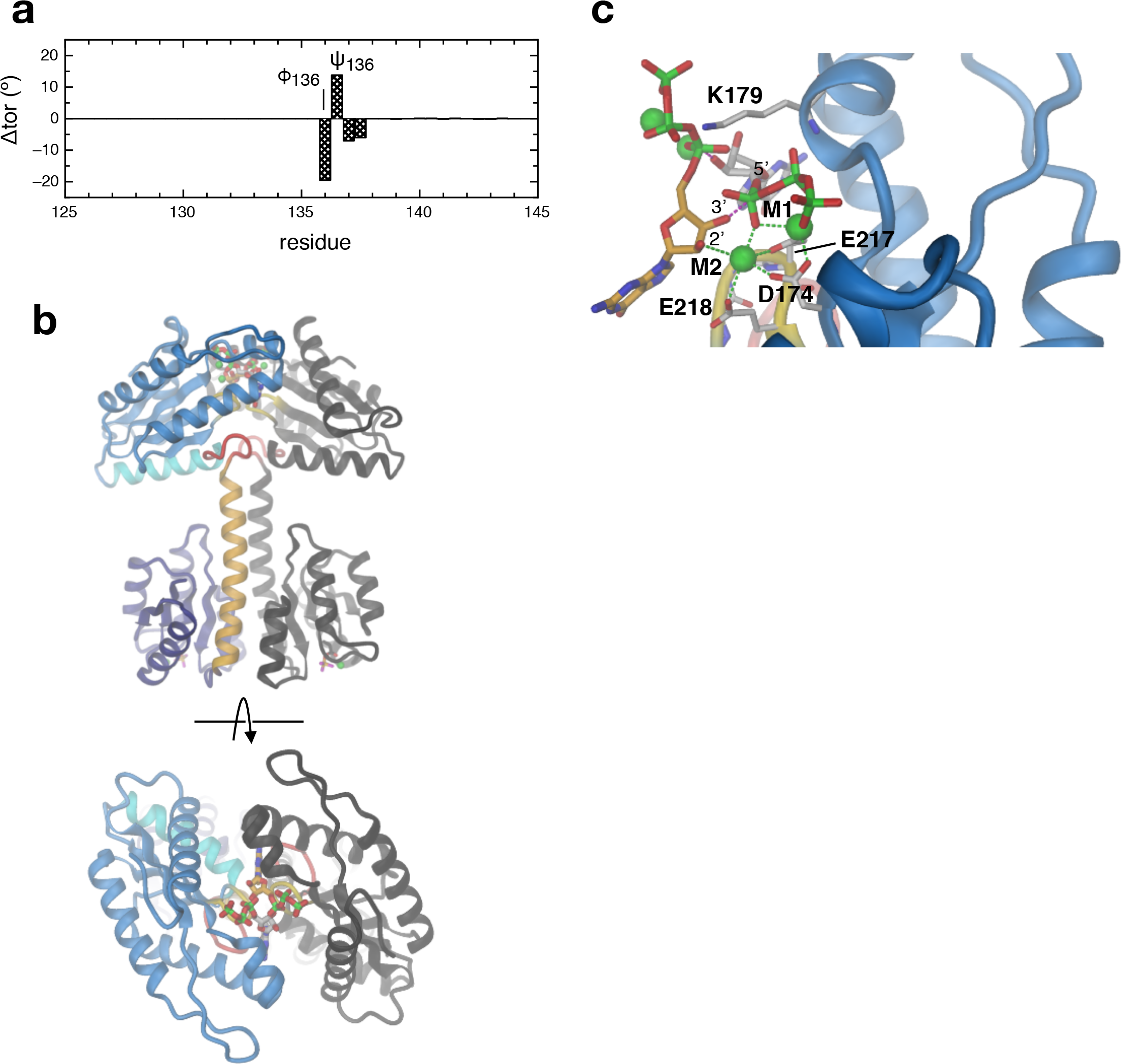
The activated Rec dimer allows formation of a catalytically competent GGDEF dimer. **a)** Changes in main-chain dihedral angles applied manually to DgcR_act to move the two GGDEF domains into catalytically competent arrangement. **b)** Model of competent DgcR (two orthogonal views) generated as described in **a**. **c)** Detailed view of the competent juxtaposition of the two GGDEF bound GTP substrate molecules. The carbon atoms of the two GTP molecules are colored in orange and gray, respectively. The O3’ hydroxyl of each ligand is poised for nucleophilic attack on the α-phosphorous (PA) of the other ligand being roughly inline with the scissile PA-O3A bond.

The details of the Michaelis-Menten complex shown in Fig. 6c are consistent with the model proposed in (Schirmer, 2016) with metal M2 coordinating the 2’-hydroxyl group and K179 hovering over the α-phosphate of the incoming substrate. There is no titratable residue close to the O3’-group. Most likely, deprotonation of the hydroxyl-group proceeds via a water molecule that could be activated by the close-by metal(s) as e.g. in adenylate cyclases (Steegborn, 2014).

In the competent dimer, there are no clashes between the catalytic domains. Molecular dynamics simulations would be required to refine the model, but it appears that D183 and D282 may interact with Y286 and H187, respectively. All these residues are conserved in diguanylate cyclases (Fig. 2 – figure supplement 1). Indeed, in the apo-structure of the constitutively active mutant of PadC (6ET7, (Gourinchas et al., 2018)) the proposed interactions seem well possible, albeit only in one half of the asymmetric structure.

There is one more conserved residue that projects to the other subunit, namely R147 (Fig. 2 – figure supplement 1). Judged by the model, it appears possible that this arginine may interact with the guanyl Hoogsteen-edge of the opposing substrate. This would be supported by a recent study on the promiscuous (accepting GTP and ATP) DGC GacA, wherein the reason for the relaxed substrate specificity was attributed to an aspartate-serine replacement of a base-binding residue (Hallberg et al., 2019). In the sub-group of promiscuous DGCs, the position homologous to R147 of DgcR is not conserved (sequence logo in Fig. 6 – figure supplement 1D of (Hallberg et al., 2019)) suggesting that the arginine is no longer important, since it cannot interact with a adenyl Hoogsteen-edge. In the same paper, the fourth residue of the GGDEF motif (equivalent to E218 in DgcR) was proposed to deprotonate the 3’-hydroxyl group of the substrate bound to the other subunit. In our model (Fig. 6c), this residue is coordinating metal M2 and clearly not close to this substrate hydroxyl group.

### Structural coupling of Rec modification with competent dimer formation

The knowledge of the structures of full-length DgcR both in its native and activated form and the model of the Michaelis-Menten complex allows to discuss in detail how signal perception (phosphorylation) is coupled to output activation in a prototypic response regulator with enzymatic function. In figure 7, this process is dissected into 5 notional steps.

**Figure 7.**
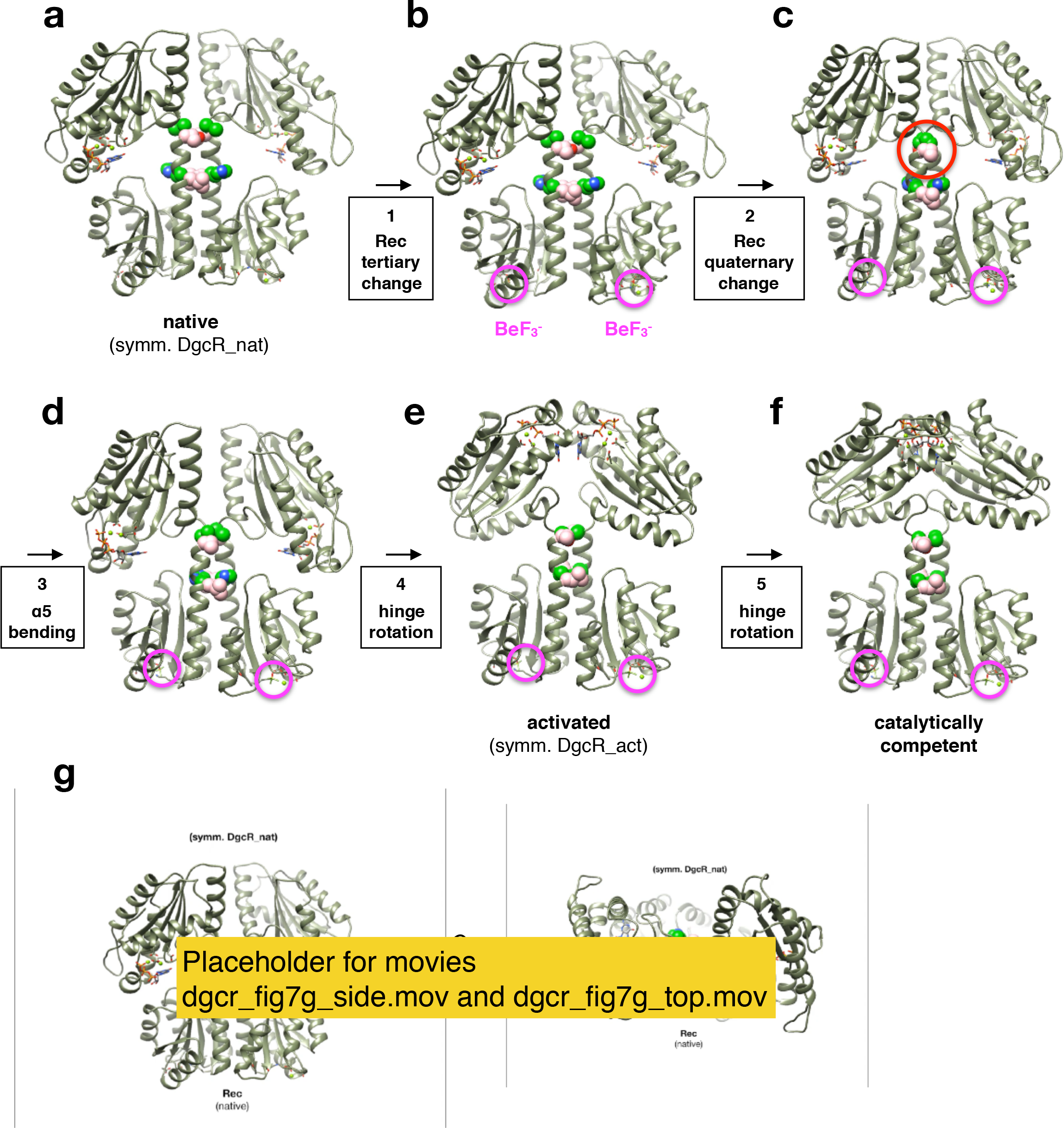
Structural transitions in DgcR upon Rec pseudo-phosphorylation (a - d) followed by GGDEF hinge motions (e-f) to attain the catalytically competent state. The structures are represented as in Figs. 1b,c, but with the residues of the conditional coiled-coil contacts shown as CPK models (residues in *d* and *e* position are shown is pink and green). The beryllofluoride moieties of the dimer are highlighted by magenta circles. **a)** DgcR_nat, symmetrized version with both GGDEF domains in B-chain orientation (cf. with Fig. 1b). **b)** As in **a**, but with beryllofluoride-induced tertiary change applied to Rec rigid_body 1 (see Fig. 3). **c)** As in **b**, but with quaternary change applied to Rec domains. Note the clash between the C-terminal ends of the coiled-coil (red circle). **d)** As in **c**, but with Rec dimer as found in symmetrized version of DgcR_act. **e)** Symmetrized version of DgcR_act (cf. with Fig. 1c). **f)** Model of catalytically competent DgcR as in Fig. 6b. **g)** Animation (two views) of the structural transitions between the states shown in panels a to f obtained by morphing. The magenta broken lines (top view, right) connect the reacting atoms of the two substrates, i.e. O3’ with Pα of the opposing substrate.

(1) Starting with a symmetrized version of DgcR_nat (Fig. 7a), aspartate pseudo-phosphorylation induces a rigid-body motion within each Rec domain (Fig. 7b, tertiary change). With an unchanged coiled-coil packing, the intermolecular α4 - α5 contacts would break up. (2) This is counteracted by a repacking of the two Rec domains (Fig. 7c, quaternary change. (3) The clashing of the C-terminal ends of the coiled-coil is relieved by slight outward bending of the helices (Fig. 7d). Obviously, these first three steps, which describe the transition of the native to the activated Rec stalk, will be tightly coupled.

The following steps invoke no direct Rec - GGDEF communication, but only an unrestricted rotation of the GGDEF domains around the inter-domain hinges. With the Rec stalk in its activated constellation, the hinges are positioned such that the GGDEF domains can attain (4) a constellation as in DgcR_act (Fig. 7e) and, finally, assemble to form (5) the catalytically competent constellation Michaelis-Menten complex (Fig. 7f). An animation of the entire structural transition from native to competent DgcR is shown in Fig. 7g.

The aspect of conformational sampling and its dependence on the coiled-coil register and the dynamics of the entire enzyme has been discussed before for PadC (Gourinchas et al., 2018). Whether the asymmetric GGDEF dimer obtained for mutated PadC is of functional importance needs further studies. Although such a state would probably be compatible with our model, it is not mandatory for the proposed mechanism in which the two phosphodiester bonds could be formed quasi-simultaneously. Furthermore, we suggest that the competent GGDEF dimer would assemble autonomously due to electrostatic and steric complementarity, in particular in presence of the substrates that interact with K179 and M2 of the opposing domain (Fig. 6c), thus not requiring any direct interaction between input and output domains.

### Allosteric inhibition by product mediated domain cross-linking

Allosteric product inhibition by c-di-GMP is a well-known feature of many DGCs (Christen et al., 2005)(Wassmann et al., 2007)(Schirmer, 2016)(Chou and Galperin, 2016). Hereby, dimeric c-di-GMP mutually cross-links a RxxD motif (primary I-site, I_p_) on one GGDEF domain with a secondary I-site (I_s_) on the other GGDEF domain and vice-versa. The crystal structure of DgcR obtained in presence of c-di-GMP (DgcR_inh) was determined to 3.3 Å by molecular replacement and is shown in Fig. 8a. There are three symmetric dimers in the asymmetric unit. Each dimer shows a Rec stalk in native conformation and the two GGDEF domains have their active sites facing outwards. Dimeric c-di-GMP is cross-linking the domains by interacting with the R206xxD209 motif of one subunit and R163* from α0’ of the other subunit (Fig. 8b). Due to symmetry, there are two isologous crosslinks within the DGC dimer. Comparison with the PleD/c-di-GMP complex (Wassmann et al., 2007) (Fig. 8c) shows very similar binding, but with PleD providing an additional arginine (R390) to the I_p_-site. The DgcR equivalent (R237) is too distant to interact, but this may happen with the Rec stalk in the activated conformation. Arginines 163* (DgcR) and R313* (PleD) fulfil the same role in c-di-GMP binding, but are not homologous on the sequence level. Indeed, it has been noted earlier that among GGDEF sequences (Paul et al. 2007) arginines are enriched at either position. The unique c-di-GMP stabilized GGDEF arrangement that differs drastically from DgcR_nat (Fig. 1b) again demonstrates the large flexibility provided by the inter-domain hinge.

**Figure 8.**
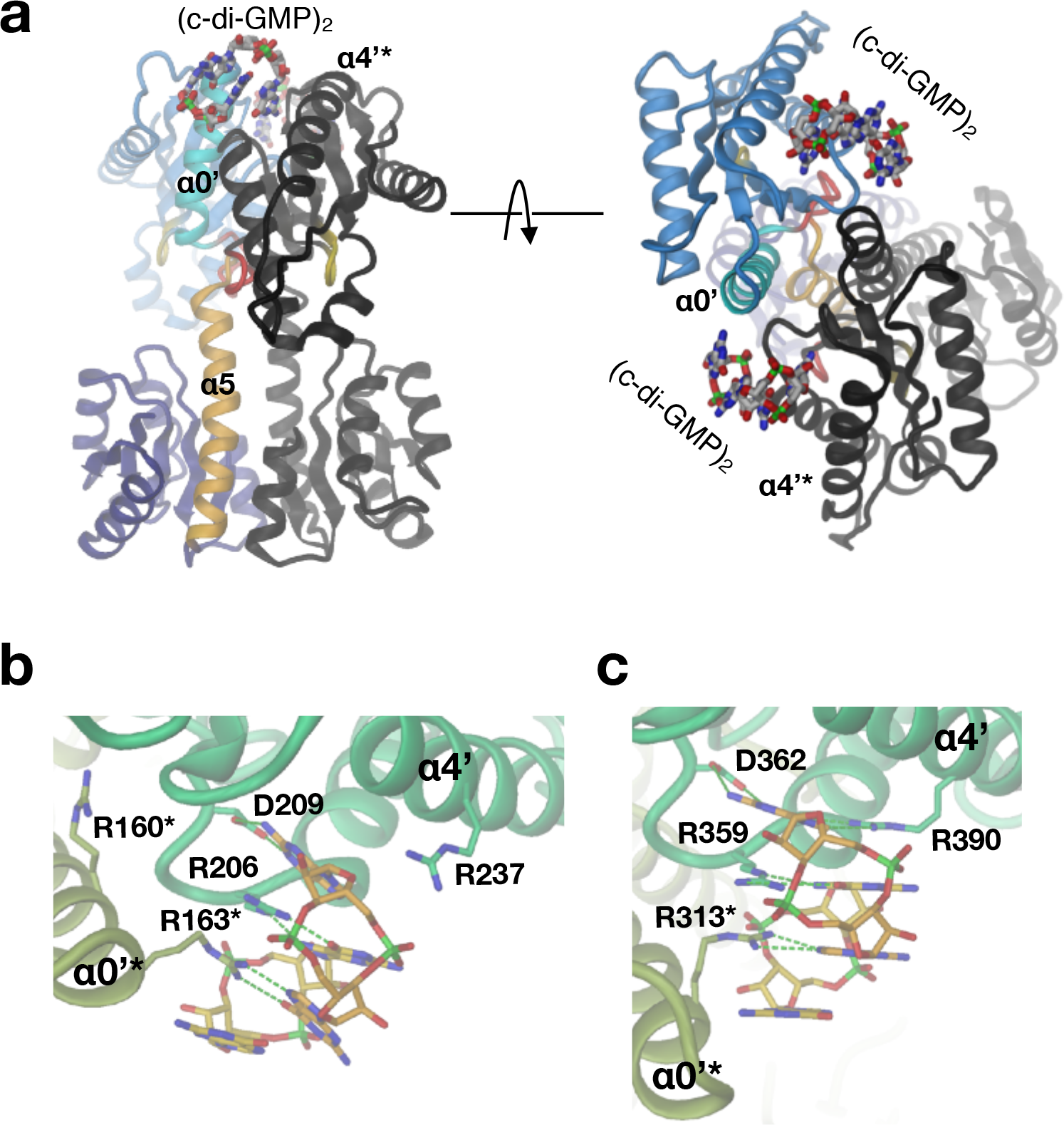
Dimeric c-di-GMP cross-links the GGDEF domain of the dimer. **a)** Side and top views of the DgcR/c-di-GMP complex (DgcR_inh). Representation as in Fig. 1b. **b-c)** Detailed comparison of the c-di-GMP binding mode in DgcR (**b)** and PleD (2V0N) (**c**). Protomers are distinguished by color hue. H-bonds are shown as green broken lines.

### Kinetic analysis of DgcR activity reveals delay in non-competitive feed-back inhibition

The effect of activation and I_p_-site mutation on DgcR catalysed c-di-GMP production was studied by a real-time nucleotide quantification assay (online ion-exchange chromatography, oIEC, Agustoni et al., in preparation, see Methods). Figure 9a shows that product formation catalysed by native wild-type DgcR (wt) gradually decreases early-on (when there is still a large excess of substrate), indicative of non-competitive product inhibition. Indeed, the progress curve was found consistent with the respective classical model with a low k_cat_ of about 0.01 s-1 and a relatively large K_i_ of about 30 μM (Tab. 2).

**Figure 9.**
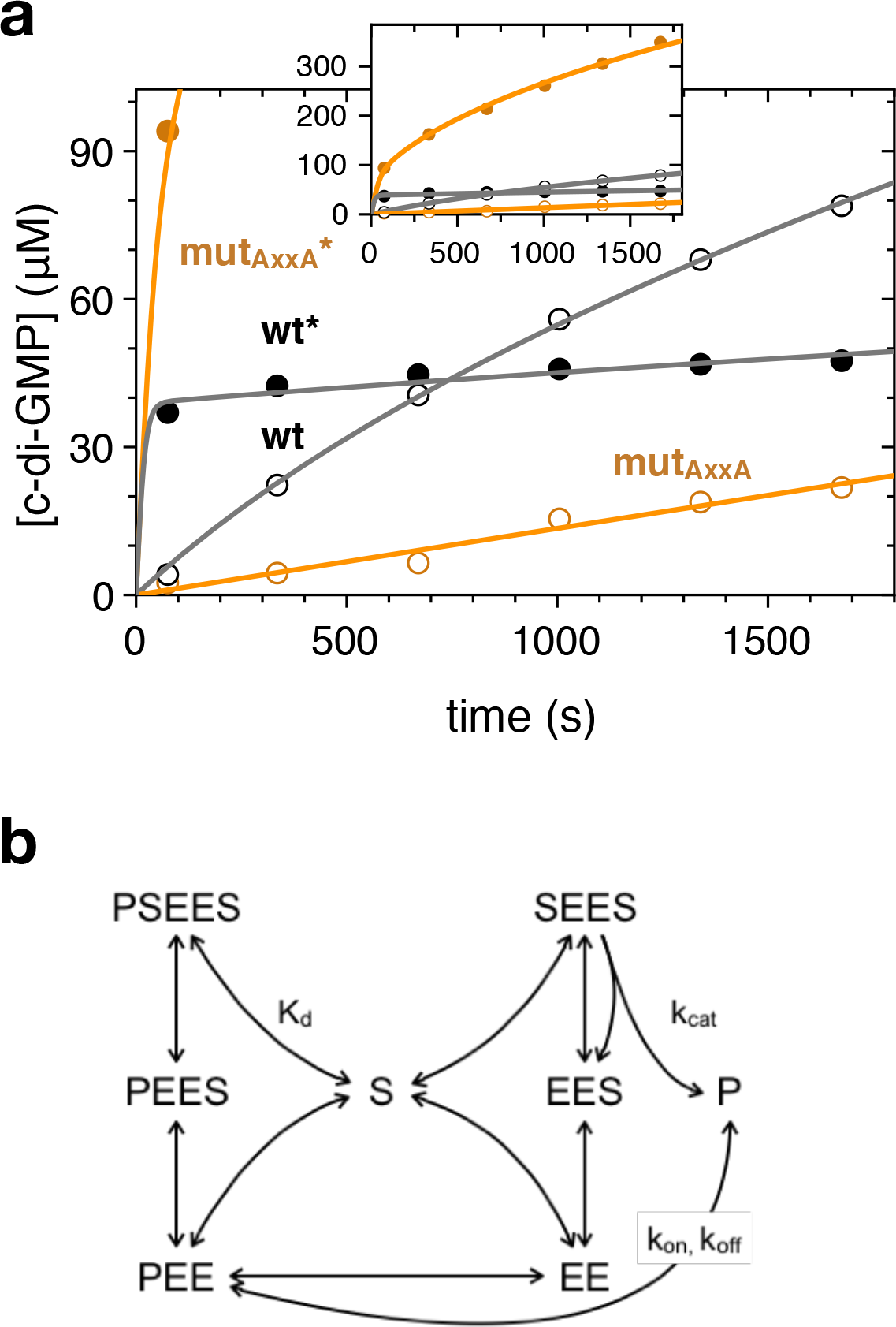
Enzyme kinetics of DgcR. **a)** Enzymatic progress curves of DgcR and DgcR_AxxA_ in the native and in the activated (indicated by an asterisk) state. Experiments were performed at 5 μM enzyme and 500 μM GTP substrate concentration. Symbols denote oiEC measurements, continuous lines represent fit of the kinetic model shown in panel **b** to the data with parameters listed in Table 2. **b)** Kinetic model of diguanylate cyclase activity controlled by non-competitive product binding. Substrate (S) binding to the dimeric enzyme (EE) is modeled with the equilibrium dissociation constant K_d_ and assumed to be unaffected by the presence of S in the second binding site or of product (P) in the allosteric site. Product binding is modeled kinetically with rate constants k_on_ and k_off_. Note that the model considers simply one instead of four product binding sites on the enzyme. Only the Michaelis-Menten complex with two bound substrate molecules and no bound product (SEES) is competent to catalyze the S + S → P condensation reaction (with turn-over number k_cat_).

**Table 1.** Data collection and refinement statistics.

|  | DgcR_nat | DgcR_act | DgcR_inh |
| --- | --- | --- | --- |
| **Data collection** |  |  |  |
| Synchrotron source | SLS, PXIII | SLS, PXI | SLS, PXIII |
| Resolution | 41 - 2.2 (2.3 - 2.2) | 41 - 2.8 (2.9 - 2.8) | 48 - 3.3 (3.4 - 3.3) |
| Space group | C 2 | P 2<sub>1</sub> 2<sub>1</sub> 2 | P 2<sub>1</sub> |
| a, b, c (Å) | 137, 39, 146 | 133, 247, 41 | 123, 73, 126 |
| $\alpha$, $\beta$, $\gamma$ (°) | 90, 109, 90 | 90, 90, 90 | 90, 118, 90 |
| Total reflections | 191690 (18770) | 205975 (19551) | 111858 (9780) |
| Unique reflections | 38087 (3727) | 34912 (3320) | 29577 (2768) |
| Multiplicity | 5.0 (5.0) | 5.9 (5.7) | 3.8 (3.5) |
| Completeness (%) | 98 (98) | 94 (97) | 99 (94) |
| Mean I/sigma(I) | 10.7 (1.2) | 8.6 (1.5) | 6.7 (1.6) |
| R-merge | 0.09 (1.20) | 0.29 (1.13) | 0.17 (0.75) |
| R-pim | 0.05 (0.60) | 0.13 (0.51) | 0.10 (0.46) |
| CC1/2 | 0.99 (0.70) | 0.90 (0.60) | 0.99 (0.70) |
| **Refinement** |  |  |  |
| R-work/R-free (%) | 23.1/25.7 | 24.8/30.1 | 25.7/31.4 |
| Number of molecules/a.u | 2 | 4 | 6 |
| Number of atoms | 4804 | 9545 | 14641 |
| protein | 4672 | 9312 | 14070 |
| ligands | 101 | 184 | 552 |
| water | 31 | 49 | 19 |
| RMS(bonds) | 0.008 | 0.009 | 0.023 |
| RMS(angles) | 1.45 | 1.52 | 3.96 |
| Ramachandran favored (%) | 97.08 | 95.14 | 95.19 |
| Ramachandran allowed (%) | 2.23 | 4.86 | 3.89 |
| Ramachandran outliers (%) | 0.69 | 0 | 0.92 |
| Average B-factor (Å<sup>2</sup>) | 65.91 | 59.80 | 81.37 |
| protein | 66.24 | 59.87 | 82.06 |
| ligands | 56.65 | 57.08 | 64.83 |
| water | 46.87 | 55.74 | 55.87 |
| PDB code | 6ZXB | 6ZXC | 6ZXM |

Statistics for the highest-resolution shell are shown in parentheses.

**Table 2.** Kinetic parameters of DgcR diguanylate cyclase activity

|  | **K<sub>d</sub> (M) +** | **k<sub>cat</sub> (s<sup>-1</sup>)** | **inhibition type** | **k<sub>on</sub>**<br>**(10<sup>3</sup>/M<sup>-1</sup>·s<sup>-1</sup>)** | **K<sub>i</sub> (M)** |
| --- | --- | --- | --- | --- | --- |
| **DgcR** | 1.0·10<sup>-5</sup> | 0.0089 | non-competitive,<br>equilibrium | n/a | 2.8·10<sup>-5</sup> |
| **DgcR\*** | 1.0·10<sup>-5</sup> | 0.2 ++ | non-competitive, slow | 5.2 | 6.2·10<sup>-8</sup> |
| **mut_AxxA** | 1.0·10<sup>-5</sup> | 0.0015 | none | n/a | - |
| **mut_AxxA\*** | 1.0·10<sup>-5</sup> | 0.2 ++ | non-competitive, slow | 7.9 | 1.1·10<sup>-5</sup> |

Parameters derived from Fig. 9a.
+ represents lower limit as obtained from simulations
++ represents upper limit as obtained from simulations

A very different behaviour was observed for activated DgcR (wt*) that produced very quickly (< 75 s) a substantial amount of product yielding a lower boundary for k_cat_ of 0.1 s-1 (Fig. 9a). This was followed by a phase of very small, virtually constant velocity. Such phenotype was clearly inconsistent with classical equilibrium models and seemed indicative of a slow transition to the product-inhibited state. Mechanistically, this transition would comprise (fast) product binding and (slow) re-organization of the two GGDEF domains to acquire the inactive product cross-linked configuration (Fig. 8).

The progress curves were fitted with the kinetic model shown in Fig. 9b. Independent binding of two substrate molecules (S) to the dimeric enzyme (EE) was parametrised with an equilibrium constant K_d_ (assuming fast substrate binding), whereas the transition between active and inactive states was modelled kinetically with an effective second-order rate constant k_on_ (dependent on product and enzyme concentration) and a first-order rate constant k_off_ with the inhibitory constant given by K_i_ = k_off_/k_on_. Note that for simplicity the model considers only one product binding site on the dimeric enzyme, while there are actually four (two c-di-GMP dimers). This simplification will affect the nominal value of K_i_. Full kinetic modelling without this simplification and with explicit modelling of the conformational enzyme transition has been postponed to a follow-up study.

The kinetic model fits the biphasic curve of wt* very well (Fig. 9a) yielding the parameters given in Table 2. The k_cat_ of 0.2 s-1 together with the slow kinetics of the active to inactive transition explains the large build-up of product in the initial phase, which is followed by very low residual activity of the (equilibrated) sample due to the low K_i_ of 62 nM.

To validate the involvement of the RxxD motif in feed-back product inhibition as suggested by the crystal structure (Fig. 8) and shown for many other DGCs, but also to scrutinise the kinetic model, the motif was mutagenised to AxxA. Indeed, the activated mutant (mut_AxxA_*) was found to be drastically, though not fully, inhibition relieved (Fig. 9a, inset). The curve is consistent with an unchanged k_cat_, but a drastically (almost 200-fold) increased K_i_ as compared to wild-type (Tab. 2). Apparently, the mutations did not completely abolish inhibition with the remaining residues of the primary and secondary I-site possibly still enabling (weak) product binding (Fig. 8).

In contrast to the native wild-type enzyme (wt), the activity of the native mutant (mut_AxxA_) was lower, which may be due to a detrimental, long-range effect of the mutations on the active site geometry. Notably, there was no indication of product inhibition. Thus, for both wild-type and mutant DgcR the activated state is more susceptible to product inhibition. Whether this is due a sub-optimal product mediated backside cross-linking in the native state as suggested by the crystal structure (Fig. 8) has to await further structural investigations.

Summarizing, activated DgcR shows a pronounced initial burst of activity before entering the product inhibited state with a rather slow kinetics probably reflecting domain reorganization. The kinetic model (Fig. 9b) proved to reproduce all measured progress curves and the parameters (Tab. 2) reflect the impact of activation and I_p_-site mutation.

### Rec - GGDEF linker sequence profiles are consistent with register shift mechanism

DgcR has been selected as a prototypic Rec - GGDEF enzyme of relatively small size (298 residues), but bioinformatic analysis showed that the linker length can vary considerably in this class of DGCs. This was surprising considering that the linker has a defined structure and seems crucial for signal transduction. However, the linker length histogram (Fig. 10a) shows that the lengths are not distributed uniformly, but exhibit discrete values separated by multiples of 7 (groups 1 to 6, with DgcR and WspR belonging to groups 1 and 4, respectively). Thus, members of the groups would merely differ in the number of double helical turns when forming parallel coiled-coils. Indeed, the individual sequence logos can easily be aligned (Fig. 10b) to reveal the striking repeat of leucines in every 7^th^ position (heptad position *a*). Most interestingly, the last (and to a lesser degree the last but one) heptad repeat at the C-terminal end (Fig. 10c) shows a conserved *axxdexx* pattern as in DgcrR (Fig. 5). Thus, a common binary register shift mechanism seems likely for members of all the groups. Group 0 (Figs. 10a, b) does not obey the linker length rule. Since it also has an (S/N)PLT instead of a DxLT motif, it probably has a different linkage and, therefore, activation mechanism.

**Figure 10.**
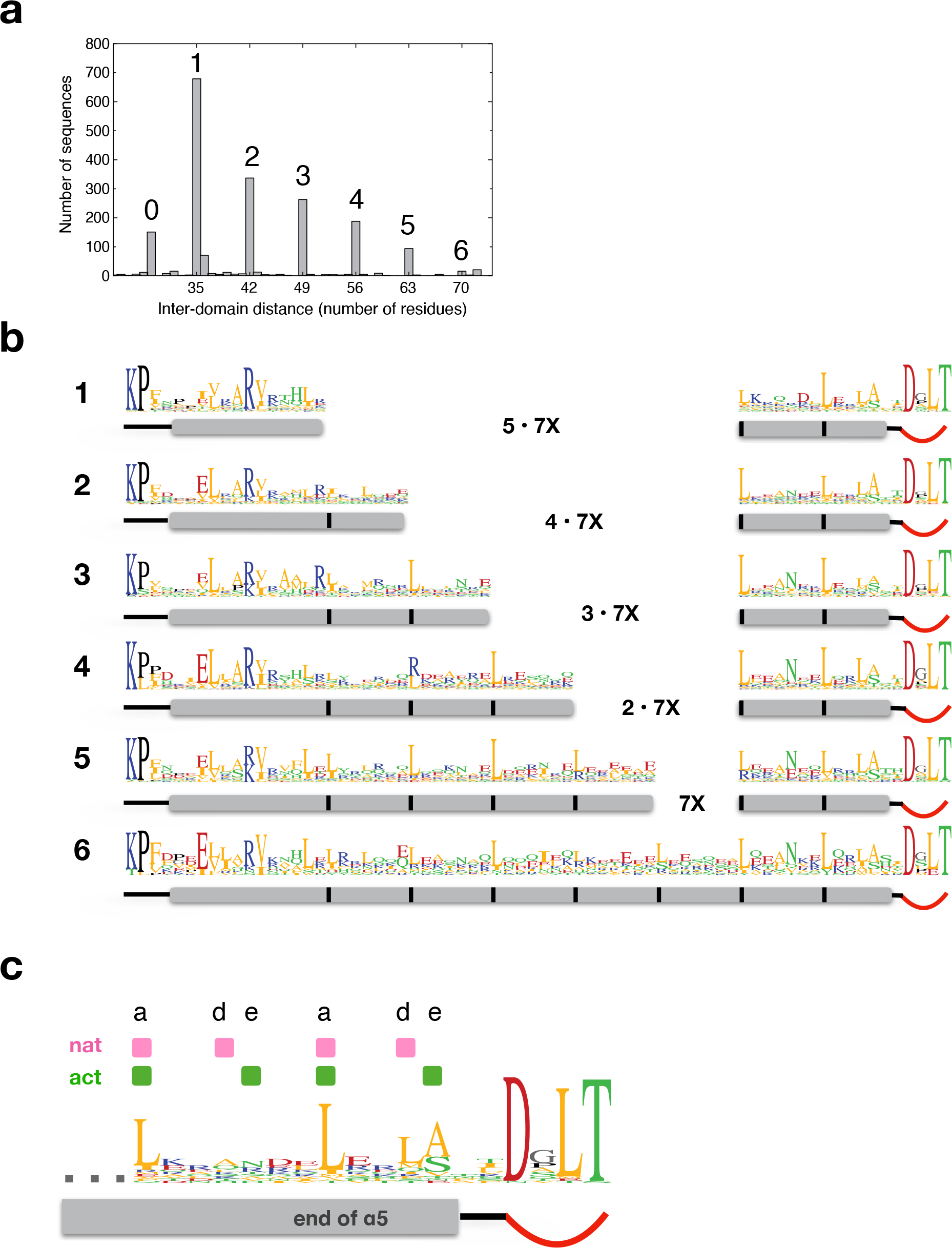
The linker helices of Rec-GGDEF proteins show heptad repeat patterns and discretized lengths. **a)** Histogram of inter-domain distances as measured from Rec KP motif to GGDEF DxLT motif. **b)** Sequence logos of inter-domain linkers of groups 1 to 6 of panel **a**. DgcR belongs to group 1. The grey rectangles symbolise the predicted α5 helices, black bars indicate recurring hydrophobic positions spaced with a distance of 7. Data were compiled from 1991 Rec-GGDEF sequences, see Methods. **c)** Overall logo of C-terminal part of all sequences shown in panel **b**. Positions engaged in parallel coiled-coil interactions in DgcR_nat and DgcR_act, are indicated in pink and green, respectively (see also Fig. 5b).

A similar pattern of discretized coiled-coil lengths has been reported for PAS – GGDEF and LOV - GGDEF proteins (Möglich et al., 2009) (Glantz et al., 2016), which makes it tempting to speculate that input to effector signal transduction might work similarly as in Rec - GGDEF enzymes. However, further investigations into their sequence profiles are needed to see whether they also exhibit ambiguous *axxdexx* heptad repeats.

## Conclusion

The presence of coiled-coil linkers between N-terminal regulatory and catalytic GGDEF domains in many diguanylate cyclases has been described and their role in signal transduction discussed (De et al., 2009)(Glantz et al., 2016; Möglich et al., 2009; Schirmer, 2016). Changes in the crossing angle or the azimuthal orientation of the helices upon activation were anticipated, but a repacking of the interface was not discussed, which was then seen first in the comparison of inactive and a constitutively activated mutant of light-regulated PadC (Gourinchas et al., 2018). The now presented detailed structural analysis of DgcR in its native and pseudo-phosphorylated form allowed a comprehensive dissection of the activation process for a full-length, wild-type Rec - GGDEF enzyme (Fig. 7). Tertiary and quaternary changes in the Rec input domains lead to a register shift in the coiled-coil linker repositioning the inter-domain hinge and, thus, the propensity of the GGDEF domains to attain the catalytically competent dimer constellation.

A register shift in the coiled-coil linker may be operational also for other enzymes with predicted coiled-coil linkers, e.g. DGCs with N-terminal GAF domains or trans-membrane helices. LOV sensor domains that carry a flavin-nucleotide chromophore and have been studied very well as part of HKs (Glantz et al., 2016)(Möglich, 2019) are different in that the coiled-coil forming Jα helix is not part of the core fold, but rather an extension of the C-terminal Iβ-strand that projects outward in the same direction. It has been shown that, upon light activation, the two Iβ - Jα junctions of the dimer increase their distance considerably (Röllen et al., 2016) probably causing a change in the crossing angle and/or the super-twist of the Jα coiled-coil in the full-length protein to control activity as discussed in the recent review by Möglich (Möglich, 2019). Most likely, GAF domain proteins control GGDEF activity in a similar way, due to the structural similarity with LOV including the predicted C-terminal coiled-coil (Möglich et al., 2009). HAMP domains have been shown to operate as rotary switches (Hulko et al., 2006). How such a change will affect the geometry of the C-terminal coiled-coil in respective DGCs has not been studied, but it will surely affect the relative disposition of the hinges that lead to the catalytic domains and, thus, activity.

Apparently, the coiled-coil linker is a versatile and effective means of transmitting a signal between domains without requiring direct interactions between them, which, obviously, is of paramount advantage for their modular combination in evolution. The same principle seems to apply also for HKs, many of them both are controlled by the same kind of input domains as DGCs and exhibit a coiled-coil preceding the DHp α1 bundle (Diensthuber et al., 2013) (Wang et al., 2013). Bioinformatic analyses (Anantharaman et al., 2006)(Glantz et al., 2016) may now be extended to test for the occurrence of “slippery” heptad repeats in coiled-coil proteins in general to reveal proteins potentially signaling *via* a coiled-coil register shift.

## Material and methods

### Protein expression and purification

*E. coli* BL21 (DE3) cells transformed with pET-28a vector containing DgcR full-length construct purchased from Genescript Inc. were incubated at 37 °C with agitation until they reached the optical density of 0.8 - 0.9. Expression was then induced by the addition of IPTG (Isopropyl β-D-1-thiogalactopyranoside) at a final concentration of 400 μM for 4 hours at 30 °C. The cells were harvested after centrifugation and resuspended in a buffer composed by 20 mM Tris pH 8.0, 500 mM NaCl, 5 mM MgCl_2_, 5 mM 2-Mercaptoethanol and protease inhibitor (Roche). The lysis proceeded by 3 passages in a French press cell at a pressure of 1500 psi. After a centrifugation at 30000 x g for 50 minutes, the soluble fraction was loaded onto a His Trap HP 5 mL column (GE Healthcare) in 20 mM Tris pH 8.0, 500 mM NaCl, 5mM MgCl_2_, 5mM 2-Mercaptoethanol and 20 imidazole. DgcR was eluted using imidazole gradient of 20 mM to 500 mM in 15 column volumes. The fractions containing DgcR were further purified by size exclusion chromatography using a Superdex 200 26/600 column (GE Healthcare) in 20 mM Tris pH 8.0, 20 mM NaCl, 5 mM MgCl_2_ and 1 mM DTT. Protein was quantified using a NanoDrop 2000 spectrophotometer (Thermo Fisher Scientific).

### BeF_3^-^_ modification of DgcR

In order to produce BeF_3^-^_ modified DgcR, approximately 300 μM of DgcR in 20 mM Tris pH 8.0, 20 mM NaCl and 5 mM MgCl_2_ were incubated with a mixture of NaF at 10 mM and BeCl_2_ at 1 mM, final concentration. After gentle mixing to achieve a homogeneous solution, the sample was left at room temperature for at least 15 minutes. DgcR BeF_3^-^_ mix was then centrifuged at 4°C at 18.000 x g to remove a light precipitation formed during the process. Protein concentration was measured after the activation process and was found virtually unaltered.

### SEC-MALS analysis

Light scattering intensity and protein concentration were measured at elution from the column using an in-line multi-angle light-scattering and differential refractive index detectors (Wyatt Heleos 8+ and Optilab rEX). These data were used to calculate molar mass for proteins by standard methods in Astra 6 (Wyatt). Corrections for band-broadening, inter-detector delays and light-scattering detector normalisation were performed using a sample of bovine serum albumin in the experimental buffer, according to the manufacturer’s protocol. Samples were loaded (100 μL) at concentrations ranging from 0.4 to 10 mg/mL in presence of various ligands at a constant flow of 0.5 mL/min in 20 mM Tris pH 8.0, 500 mL NaCl, 5 mM MgCl_2_, 1 mM DTT.

### Crystallization

Crystallization attempts were performed using vapour diffusion method prepared in 3-drop MRC plates by Gryphon robot (Art Robbins Instruments) with DgcR (wild-type or I-site mutant AxxA) at a concentration of 10 mg/mL (280 μM) in 20 mM Tris pH 8.0, 20 mM NaCl, 5 mM MgCl_2_ and 1 mM DTT at 18 °C. For DgcR_nat crystallisation, 3’dGTP was added at a final concentration of 2 mM.

After 3 days, crystals could be observed in 0.2 M Magnesium sulfate, 20 % PEG 3350 from condition C8 of PEG/Ion HT crystallization kit (Hampton Research). DgcR_act was crystallised by the same protocol, but with BeF_3^-^_ treatment prior to the crystallisation set-up. After 7 days crystals were observed in a condition composed by 0.3 M Magnesium chloride hexahydrate, 0.3 M calcium chloride dehydrate, 1.0 M imidazole, MES monohydrate (acid), pH 6.5, EDO_P8K, 40% v/v ethylene glycol, 20 % w/v PEG 8000 present in condition A2 from Morpheus I crystallization kit (Molecular Dimensions). Crystallization of DgcR in the inhibited conformation (DgcR_inh) was achieved by the presence of 2.0 mM c-di-GTP. Crystals appeared after 5 days in 0.2 M potassium thiocyanate, 0.1 M Tris pH 7.5, 25% PEG 2000 MME, condition optimized from H11 of Index HT crystallization kit (Hampton Research). Crystals were frozen in liquid nitrogen and stored in a transport Dewar prior to data collection.

### Crystal data collection and structure determination

Data was collected at the Swiss Light Source (SLS), Villigen, Switzerland at 100 K and was processed using XDS (DgcR_nat), iMosflm (DgcR_act data and DgcR_inh) and CCP4i suite (Kabsch, 2010)(Potterton et al., 2018) (Battye et al., 2011). DgcR_nat structure was solved by molecular replacement using homologous structures generated from the Auto-Rickshaw pipeline web server (Panjikar et al., 2009). Subsequently, the DgcR_act and DgcR_inh structures were solved by molecular replacement using the Rec and GGDEF domains of DgcR_nat separately using Phaser (McCoy et al., 2007). The model was built using COOT and refinement was carried using Refmac5 (Emsley and Cowtan, 2004) (Murshudov et al., 2011). Structure figures were prepared using Dino (http://dino3d.org). Morphing was calculated using UCSF Chimera (Pettersen et al., 2004).

### Enzymatic analysis

DgcR wild type and DgcR_AxxA activity assays were performed at 5 μM in the presence of 500 μM of GTP in a reaction buffer composed of 100 mM Tris pH 8.0, 100 mM NaCl and 5 mM MgCl_2_. The reaction was started by substrate and product progress curves were acquired by a novel automatic chromatographic method, named online ion exchange chromatography (oIEC) (Agustoni, manuscript in preparation), in which aliquots (68 μL) are automatically withdrawn from the large reaction vessel (650 μL) and loaded into a Resource Q column (GE Healthcare) without the need for prior quenching of the reaction. This was followed by ammonium-sulfate (0 to 1 M, 20 mM tris, pH 8.0) gradient elution of the bound substances (enzyme, substrate, product). Peak areas corresponding to the c-di-GMP product were integrated and converted to concentrations using a scale factor obtained from calibration. Data was plotted and fitted using proFit (QuantumSoft).

To calculate theoretical progress curves, the partial differential equations corresponding to the kinetic scheme in Fig. 9b were set-up in ProFit and solved by numerical integration. Global fitting of this function using the Levenberg algorithm implemented in ProFit to the measured time courses of product and substrate concentration yielded the parameters listed in Table 2.

### Bioinformatic analysis

Rec and GGDEF domain HMM profiles were taken from Pfam (Finn et al., 2010) and used as input to an hmmsearch on the HMMER web server against the reference proteome database rp55 (E-values 0.01; hit 0.03) (Potter et al., 2018). 8016 sequences were found and filtered by size (< 360 residues) to exclude Rec - GGDEF sequences with additional domains. This procedure reduced the data size to 1991 sequences. A redundancy filter (< 80% pairwise identity) finally reduced the number of sequences to 1408. Global alignment was performed using Muscle (Madeira et al., 2019). From this alignment, the linker sequences (as defined ranging from the KP-motif in the Rec β5 - α5 loop to the DxLT motif at the beginning of the GGDEF domain) were extracted and clustered according to length by a custom-made Python script. For the major clusters, corresponding logos were generated using Geneious Prime 2020.1.2 (www.geneious.com) and manually aligned to account for the distinct linker lengths.

### Accession codes

Coordinates and structure factors of DgcR_nat, DgcR_act, and DgcR_inh have been deposited in the Protein Data Bank under the accession codes 6ZXB, 6ZXC, and 6ZXM.

## Supporting information

dgcr_fig7g_side.mov

dgcr_fig7g_top.mov

## Acknowledgments

We thank the beamline staff at the Swiss Light Source in Villigen and the Biophysics facility at the Biozentrum Basel for expert biophysical support. We thank T. Sharpe, E. Agustoni, U. Jenal, and T. Maier for critical reading of the manuscript. This work was supported by Grant 31003A-166652 of the Swiss National Science Foundation.

---

**Figure 1—figure supplement 1.**
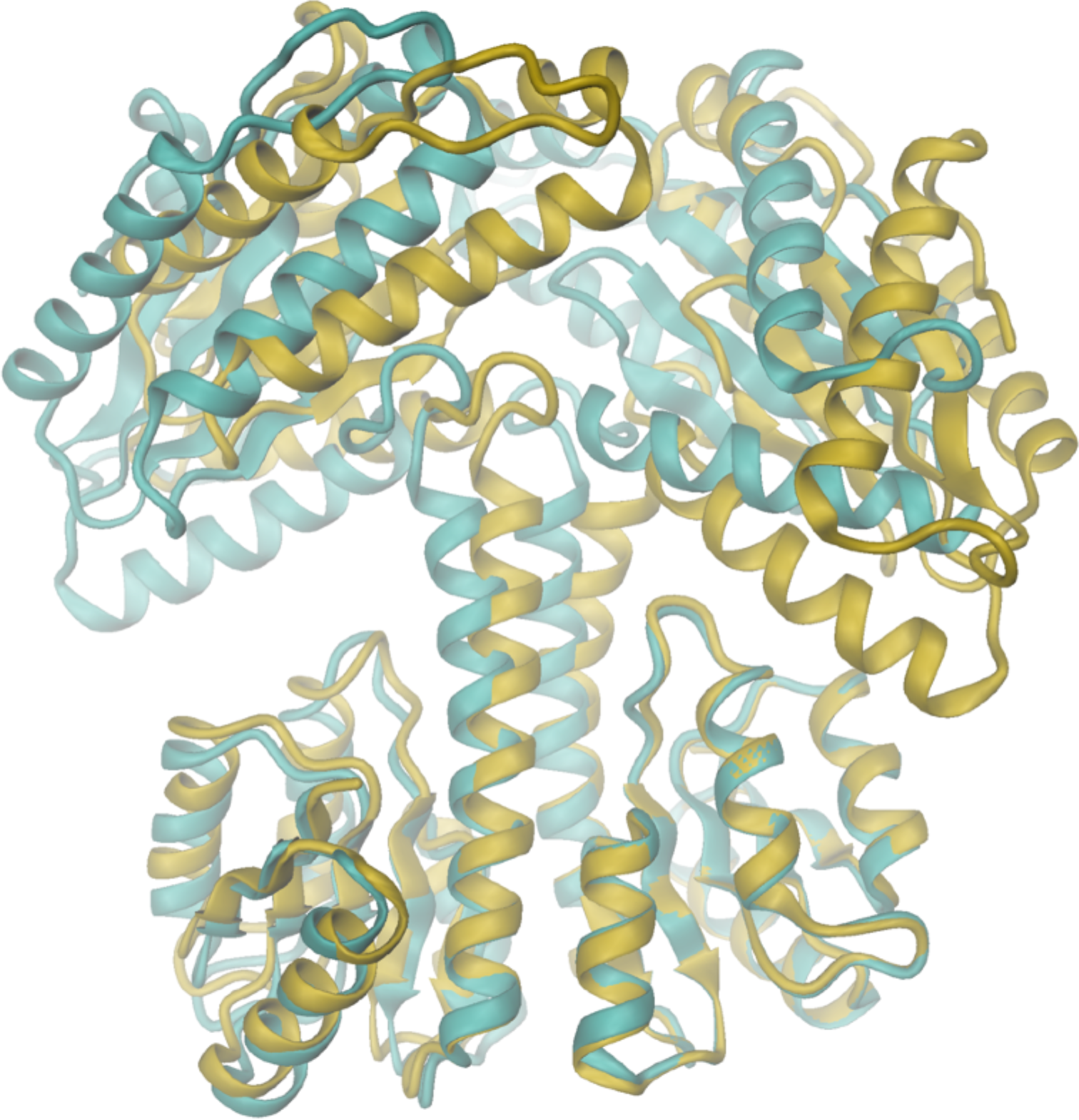
Superimposition of the two dimers of the asymmetric unit of the DgcR_act crystal structure. Dimers AB and CD were superimposed on their Rec part and are represented in turquoise and gold, respectively.

**Figure 1—figure supplement 2.**
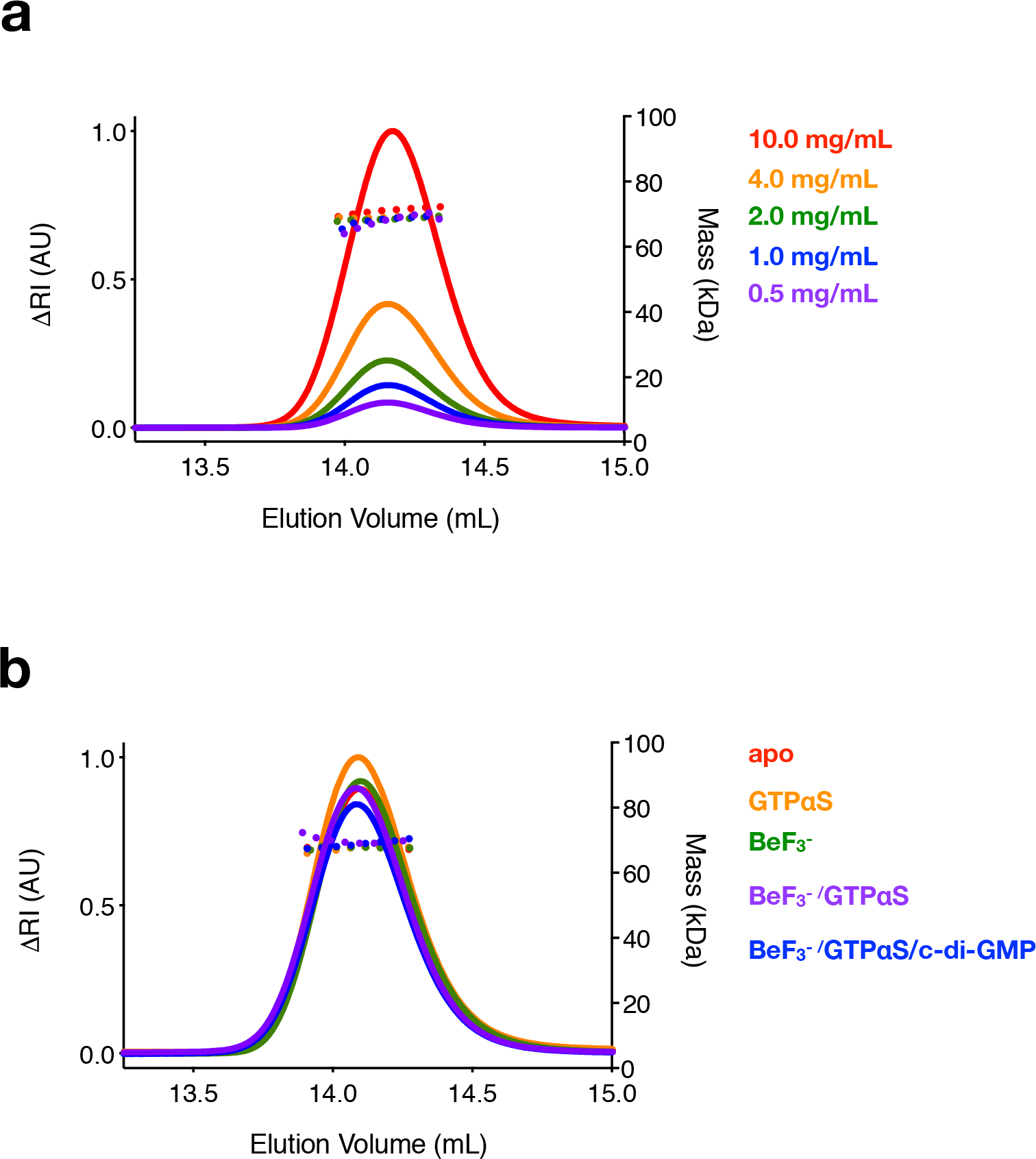
DgcR is a constitutive dimer. Oligomeric state of DgcR as analyzed by SEC-MALS as measured at the indicated **a)** loading concentrations (0.275 mM to 0.014 mM) and **b)** conditions (with GTPαS and c-di-GMP at a concentration of 2 mM and DgcR at a concentration of 0.14 mM). Molecular mass values (right axis) are shown by dotted lines and change in refractive index (left axis) by solid lines.

**Figure 2—figure supplement 1.**
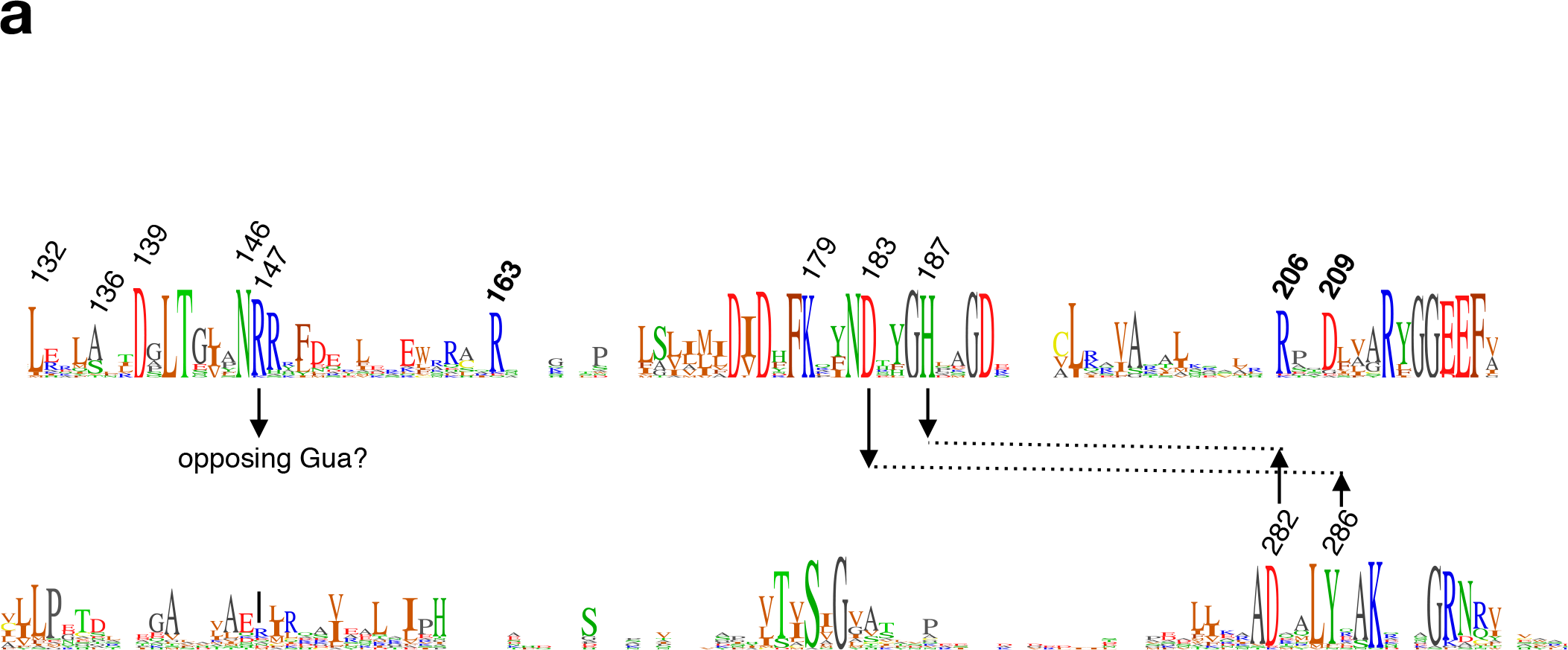
Sequence logo encompassing the GGDEF domain of Rec - GGDEF DGCs. The logo has been derived from the DgcR homologs of group 1 (see Fig. 10). Important residues are indicated by their number in DgcR (in bold for residues involved in c-di-GMP feed-back inhibition, see Fig. 8). Arrows indicate putative residues engaged in inter-domain contacts in the competent GGDEF dimer arrangement (Fig. 6b). R147 may interact with the Hogsteen-edge of the guanyl-base of the substrate bound to the opposing domain.

**Figure 5—figure supplement 1.**
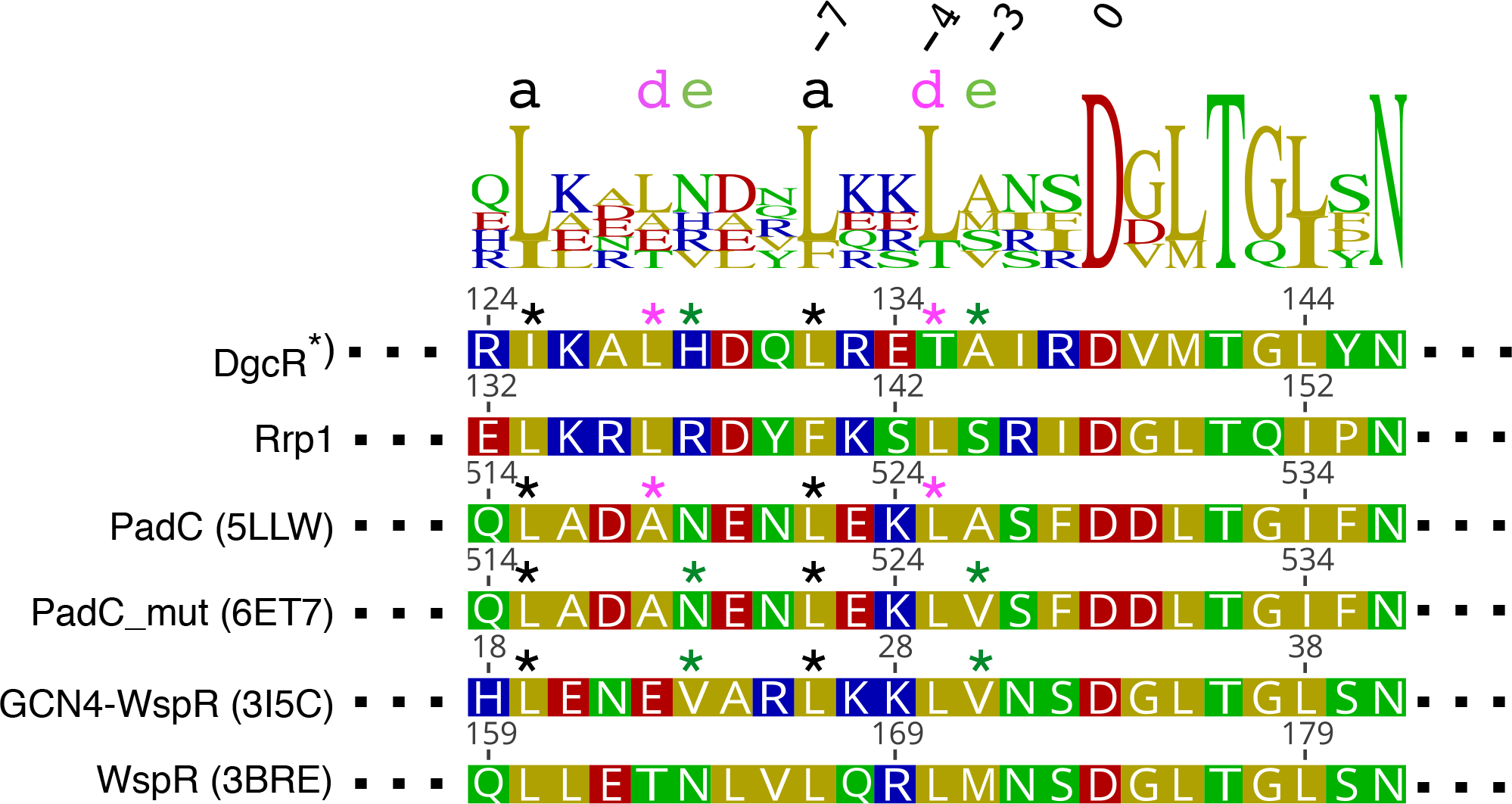
Alignment of selected DGCs comprising C-terminal end of Rec α5 and beginning of GGDEF domain. DgcR and 3 other well studied DGCs were included in the alignment: Rrp1 from *Borrelia burgdorferi*, PadC from *Idiomarina sp. A28L* and WspR from *Pseudomonas aeruginosa* (wild type and GCN4 hybrid). Sequence numbers relative to DxLT motif are given on the top. Asterisks denote crystallographically observed coiled-coil contacts (black: persistent contacts; colored: conditional contacts, i.e. contacts formed only in the native (pink) or activated (green) register). *) DgcR_nat and DgcR_act of this study.

**Figure 6—figure supplement 1.**
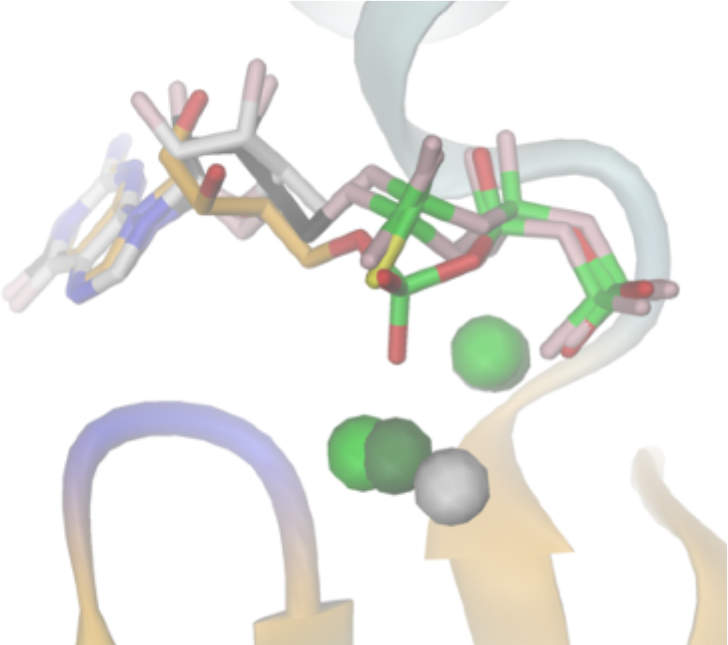
Superposition of GTP analog structures as bound to GGDEF domains. 3’dGTP/Mg^++^ as found bound to DgcR_act (cartoon) is shown with yellow carbons and light green spheres. 3’dGTP/Mg^++^ as found bound to DgcR_nat is shown with pink carbons and dark green spheres. The 3’-OH groups of 3’dGTP have been reconstructed. GTP-αS/Ca^++^ as bound to DosC (4ZVF) is shown with pink carbons and a grey sphere.

